# Surface avidity of anionic polypeptide coatings target nanoparticles to cancer-associated amino acid transporters

**DOI:** 10.1101/2025.07.28.667320

**Authors:** Ivan S. Pires, Margaret M. Billingsley, Ezra Gordon, Andrew J. Pickering, Eva Cai, Gonzalo J. Esparza, Mae L. Pryor, Alexander D. Stoneman, Darrell J. Irvine, Paula T. Hammond

## Abstract

Tumor-targeted drug delivery enhances therapeutic efficacy while minimizing toxicity. Layer-by-layer nanoparticles (LbL-NPs) coated with anionic polypeptides selectively bind to cancer cells, though the mechanisms have been unclear. Here, we integrated *in silico* and *in vitro* approaches—including gene expression analysis, receptor inhibition, and AI-based protein modeling—to show that poly(L-glutamate) (PLE)-coated LbL-NPs bind with high avidity to SLC1A5, a glutamine transporter overexpressed in cancer. We also discovered that PLE clusters SLC1A5 on the cell membrane, promoting prolonged cell surface retention. Poly(L-aspartate) (PLD)-coated NPs similarly bind SLC1A5 but also interact with faster internalizing transporters of anionic amino acids. Correlation analyses across cancer cell lines confirmed a strong link between transporter expression and nanoparticle association. These findings demonstrate that dense glutamate or aspartate presentation through electrostatically adsorbed polypeptides enables selective targeting of overexpressed transporters, providing a mechanistic framework for receptor-targeted delivery that leverages metabolic characteristics of a range of solid tumor types.

## Main

Nanoparticles (NPs) are promising vehicles for drug delivery due, in part, to their ability to modulate drug bioavailability and pharmacokinetics.^1,2^ A central challenge in cancer nanomedicine is achieving selective delivery of therapeutics to malignant cells while minimizing off-target toxicity in healthy tissues. Early strategies relied on the enhanced permeability and retention (EPR) effect, which exploits the leaky vasculature of tumors to promote NP accumulation.^3,4^ However, the EPR effect has demonstrated inconsistent efficacy across tumor types and limited clinical translation. To address these limitations, alternative models such as active transport and retention (ATR) have recently been proposed to better explain and predict NP accumulation in tumors.^5^ Beyond non-specific targeting, functionalizing NPs with antibodies, peptides, or small molecules enables more selective delivery by engaging specific cell-surface receptors.^4^ Despite its promise, this approach faces challenges, including the scarcity of truly cancer-specific targets, tumor heterogeneity, and the immunogenicity or manufacturing complexity of targeting ligands. These limitations have prompted growing interest in alternative mechanisms for tumor-specific NP binding that do not rely on conventional ligand-receptor paradigms. Among these, a promising method to modulate NP properties and enable cell-specific targeting is through the established electrostatic layer-by-layer (LbL) technique for functionalizing NP surfaces.

We previously reported that coating the surface of NPs with charged polypeptides composed of poly-L-aspartate (PLD) and poly-L-glutamate (PLE) enhanced their affinity and specificity towards cancer cells and had varied internalization rates.^6,7^ This unique targeting strategy makes LbL-NPs of interest as potentially broadly applicable cancer-targeting delivery vehicles. However, unlike some LbL-NPs with high cancer cell affinity—such as hyaluronic acid (HA)-coated particles that target CD44 receptors and promote rapid receptor-mediated endocytosis—the receptor(s) mediating specific binding of these anionic polypeptide coatings to cancer cells have remained unclear.^6^

One potential mechanism of the cancer-specific association of these polypeptide coatings is through amino acid transporters. Cancer cells often become dependent on certain amino acids such as glutamine or aspartate^8–11^, and to sustain this intracellular amino acid pool requirement, the cancer cells modulate a network of redundant transmembrane proteins that mediate cellular amino acid transport.^12^ Here, we explore the increased expression of amino acid transporters as a potential mechanism underlying the specificity of polypeptide LbL-NPs binding to cancer cells. Through a combination of experimental approaches, data analytics, and artificial intelligence protein interaction modeling, we demonstrate that LbL film assembly allows for high avidity presentation of amino acid side chains that interact with amino acid transporters overexpressed by cancer cells. Further, we show how PLE cell membrane retention is enhanced by cell surface receptor cluster formation. Lastly, we find that cellular gene expression of amino acid transporters correlates with efficient LbL-NP association while demonstrating that PLD NPs further interact with different amino acid transporters than PLE NPs, which may contribute to differences observed in their cellular trafficking. Together, these findings provide insights into the mechanisms of anionic polypeptide-based targeting for NP delivery across cancer cells.

### Polypeptide presentation and avidity regulate cancer cell binding affinity

Cancer cell-targeting LbL-NPs are formed by sequential adsorption of oppositely charged polymers onto a charged NP surface.^6,13,14^ For poly(L-glutamate) (PLE)-coated LbL-NPs, a bilayer film composed of positively-charged poly(L-arginine) (PLR) and negatively-charged PLE assembled onto a negatively-charged liposome core is sufficient to generate high affinity binding to cancer cell surfaces (**Figure 1a**).^13,15,16^ We first sought to determine if PLE has an intrinsic high affinity for cancer cells and if incorporation of this polypeptide into the LbL film coating the NP surface plays an important role in its binding activity. The ovarian cancer cell line OV2944-HM1 (HM-1) was incubated with two PLE polymers at varying degrees of polymerization (PLE_100_ or PLE_800_) or with LbL-NPs coated with PLE_100_ polypeptide (PLE_100_- NPs), and cell association was measured by flow cytometry. While increasing the polymer chain length did allow for a ∼9-fold increase in EC_50_, PLE_100_-NPs exhibited a much higher apparent affinity of binding, with an EC_50_ 50,000-fold lower than free PLE_100_, demonstrating a major effect of presentation of the PLE from NP surfaces on cell association (**Figure 1b**).

**Figure 1.**
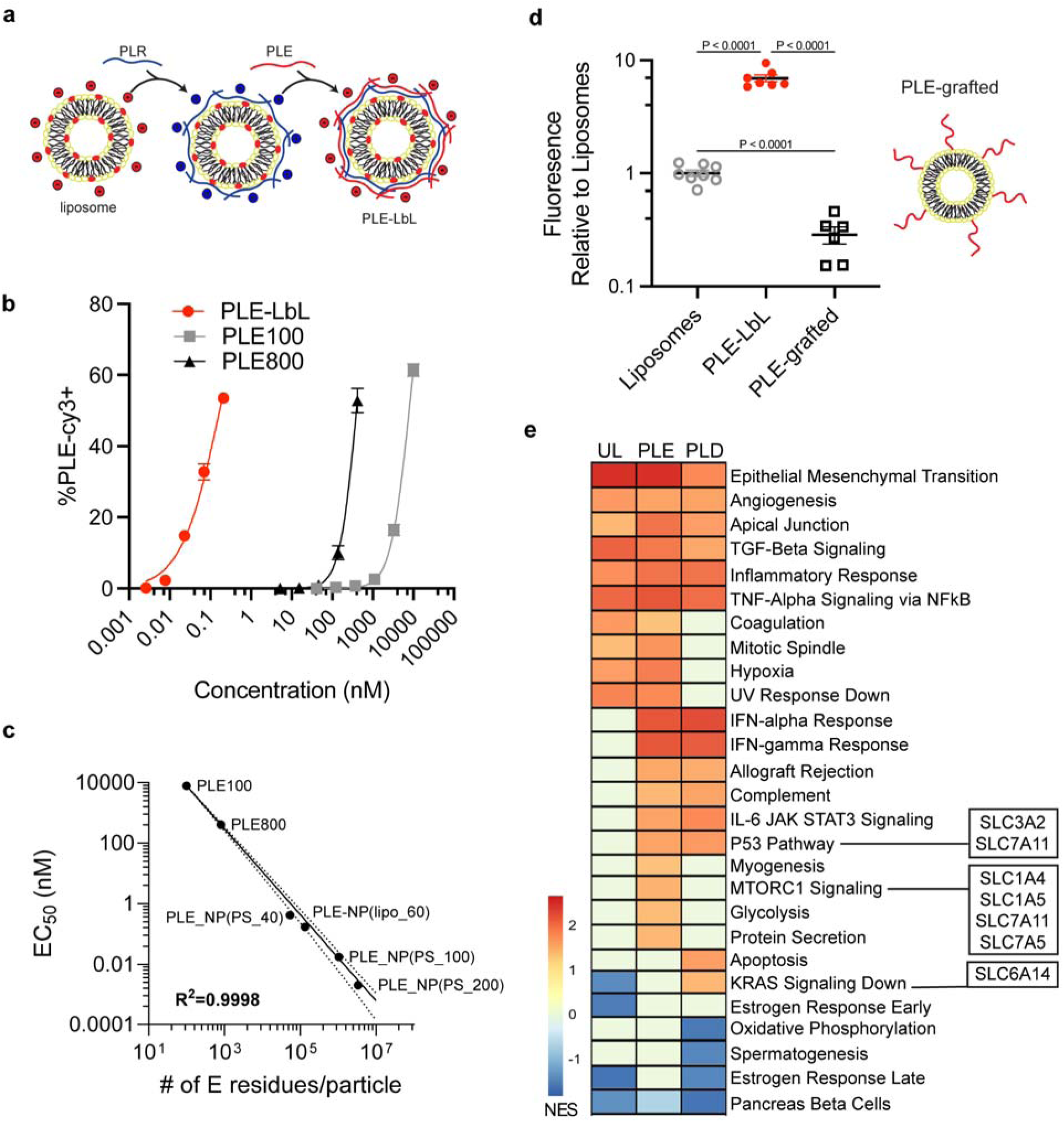
Polypeptide LbL films enable high-affinity nanoparticle binding to cancer cells. **(a)** Schematic of layer-by-layer (LbL) surface modification in which polymers of alternating charge are electrostatically layered onto the NP surface. **(b)** Fluorescently tagged PLE was dosed at varying concentrations to HM-1 cells for 4 hrs either in the LbL-film on NP (PLE_100_-NP) or as free polymers at a 100 or 800 degree of polymerization, and flow cytometry was used to measure the percentage of PLE+ cells (mean ± s.d). **(c)** Relationship between estimated number of glutamate (E) residues per particle or per molecule for free polymer and the derived EC_50_ from the experiment in (**b**) and EC_50_ of NP+ cells dosed with PLE-coated carboxylate-modified polystyrene (PS) NPs of varying diameters (40 nm, 100 nm, and 200 nm). **(d)** Fluorescently labeled NPs were dosed to HM-1 cells *in vitro* at 1 µg/mL. 4 hrs after dosing, cells were washed and NP fluorescence associated with cells was measured on a plate reader. Shown are the normalized fluorescence readings relative to an unlayered negatively charged liposome (mean ± s.d). **(e)** Data from NanoPrism^21^ was used to rank cell lines based on their association with each NP (UL, PLE, or PLD), and the gene expression of the top 100 and bottom 100 cell lines was then compared via gene set enrichment analysis (GSEA) against the hallmark gene sets. The heatmap shows the normalized enrichment score (NES) of significant gene sets (FDR q-val < 0.05) for any of the three NP groups with positive values indicating these gene sets were enriched in the cell lines that corresponded with high NP association. Genes for relevant amino acid transporters are noted next to their corresponding gene sets. Statistical comparison in **d** was performed using one-way analysis of variance (ANOVA) with Tukey’s multiple-comparisons test. Data are representative of at least two independent experiments with n = 3 technical replicates per group.

To better understand how incorporating PLE into an LbL-NP led to this increase in association, we explored the effect of glutamate residue multivalency on cancer cell affinity. We evaluated PLE_100_ and PLE_800_ free polymers, 60 nm liposomal PLE_100_-NPs, and fluorescently-labeled carboxylate-modified polystyrene (PS) NPs of varying sizes (diameters of 40, 100 and 200 nm) coated with a PLE outer layer. The varying of diameters allows us to compare a range of glutamate (E) residues per particle, and their resulting affinity towards HM-1 cells was quantified. Comparing the estimated number of glutamate (E) residues per particle (determined from maximum polymer loading based on zeta potential plateau^17^) or free polymer molecule to the observed cellular apparent affinity yielded a clear log-log relationship suggesting that the high surface avidity of glutamate residues from the LbL film drives the high-affinity interactions (**Figure 1c**).

To understand if the mode of PLE presentation from NPs is important for its high cellular affinity, we next evaluated the binding of fluorescently-labeled unmodified liposomes, liposomes coated with PLE_100_ either via an electrostatically adsorbed LbL film, or liposomes bearing PLE_100_ covalently end-grafted to the liposome surface (**Figure 1d**), and measured cell-associated fluorescence after 4 hr of in vitro incubation. We maximized the amount of grafted PLE polymers on the NPs to yield liposomes of ∼100 nm in size and similar negative surface charge (**Extended Data Fig. 1a-c**). Increasing the grafting density beyond 0.15 weight equivalents of PLE to total lipids led to disc or micelle formation due to charge and steric repulsion.^18^ While PLE-LbL coating increased NP binding ∼10-fold compared to unlayered (UL) liposomes, grafted PLE polymers unexpectedly reduced binding by a similar magnitude (**Figure 1d**). This may suggest that grafted polypeptides create extended brush-like layers that sterically inhibit rather than promote cell binding.^19,20^ Chain conformations of electrostatically layered polymers in LbL films are quite different and present as loops and trains bound on the surface.^20^ These results suggest that the nature of polypeptide presentation from the NP surface is critical for determining cellular interactions. Notably, while it is possible to generate monolayer PLE-coated NPs onto cationic liposomes (**Extended Data Fig. 1d-e**), these did not confer increased cellular association over unlayered NPs (**Extended Data Fig. 1f**) likely due to both increased non-specific association of underlying cationic NPs and low film stability of monolayer coated NPs.

Across these evaluations of cell association, the cellular affinity of PLE was highest when presented via the LbL NP platform and directly correlated with glutamate residue valency per particle. We sought to understand what transcriptional profiles promote binding of the polyvalent glutamate-presenting PLE-NPs to the surface of cancer cells by mining our previously published nanoPRISM dataset, which contains data from high throughput screens measuring the association of various NPs with or without LbL coatings to a library of 488 human cancer cell lines.^21^ Specifically, we looked to compare the performance of UL liposomes with liposomes layered with PLE. In addition to PLE LbL-NPs, we also evaluated the binding of poly-L-aspartate (PLD) coated NPs given the close similarity between these anionic polypeptides and their cancer cell targeting properties demonstrated in previous work.^6^ Within each NP group— UL, PLE, and PLD—we compared the gene expression of the 100 cell lines exhibiting the highest NP association with the gene expression of the 100 cell lines exhibiting the lowest NP association. We then rank-ordered the genes based on the highest p-value adjusted fold-difference in expression and performed Gene Set Enrichment Analysis (GSEA) against hallmark gene sets (**Figure 1e).** From this analysis, many gene signatures were identified. For example, epithelial-to-mesenchymal (EMT) transition was enriched for high association with all three NP groups which may be due to higher overall cellular activity and nanoparticle uptake. Moreover, consistent with prior experiments, STAT3 signaling was a significant hit for increasing the association of PLE-liposomes.^22^

In further analysis, we focused on identifying cell surface-expressed features that mediate the interaction between the anionic, glutamate-rich PLE coating and cancer cells. Given that PLE is composed entirely of glutamate residues, and that we observed increased cellular association with increasing glutamate presentation, we hypothesized that specific amino acid transporters could play a role in mediating this interaction. We searched for relevant amino acid transporters among the top enriched gene sets corresponding to high-association cell lines for PLE-NPs, PLD-NPs, or both. This effort identified SLC1A4, SLC1A5 and SLC7A5 for PLE-enriched gene sets, SLC6A14 for PLD-enriched gene sets, and SLC3A2 and SLC7A11 for both PLE- and PLD-NPs enriched gene sets (**Figure 1e**). Notably, all the identified genes are direct transporters or promote glutamine uptake with highly interconnected functions. For example, SLC3A2 and SLC7A5 encode for proteins that heterodimerize to achieve functional expression in the plasma membrane, SLC1A5 and SLC7A5 often act in opposition to balance intracellular glutamine concentration, and SLC7A11 increases SLC1A5 glutamine uptake by exporting intracellular glutamate.^23–25^ Additionally, a number of these transporters are commonly overexpressed in cancer cells including SLC1A5, SLC7A5, and SLC7A11.^26–28^ These results suggested an influence of these different amino acid transporters—especially glutamine transporters—on LbL- NP/cell association.

### Glutamine transport inhibitors block PLE-NP binding to cancer cells

Given the potential for polypeptide coatings to interact with amino acid transporters, we next evaluated whether amino acid transport inhibitors could influence LbL-NP binding. We focused on PLE-coated LbL-NPs as our model system based on its unique cell membrane retention property compared to other NPs and promising preclinical utility for therapeutic targeting of ovarian cancer and glioblastoma.^6,7,13,15,16,29^ HM-1 cells were pretreated with various concentrations of the glutamine uptake inhibitors L-γ-Glutamyl-p-nitroanilide (GPNA) or V9302. We then added fluorescently labeled LbL-NPs to the cells—comparing UL- and PLE- NPs with NPs layered with HA, PAA, or dextran-sulfate (DXS). We utilized these controls as HA-NPs are known to bind CD44, PAA-NPs have reduced cellular association, and DXS-NPs are known to associate with immune cells over cancer cells^30,31^. Across these LbL-NP groups, both glutamine uptake inhibitors enabled a significant decrease in cell association only for PLE- NPs (**Figure 2a-c**), which supports the identification of glutamine transporters as binders for PLE-NPs. Consistent with its ∼100-fold greater potency in glutamine uptake inhibition relative to GPNA^32^, V9302 exhibited a significantly stronger effect on PLE-NP binding. Further, both glutamine uptake inhibitors were able to demonstrate dose-dependent blocking of PLE-NPs while TFB-TBOA—a potent aspartate and glutamate uptake inhibitor^33^—did not affect PLE-NP binding even at concentrations 1000x higher than its reported IC_50_ (∼10-100 nM) (**Figure 2a**).

**Figure 2.**
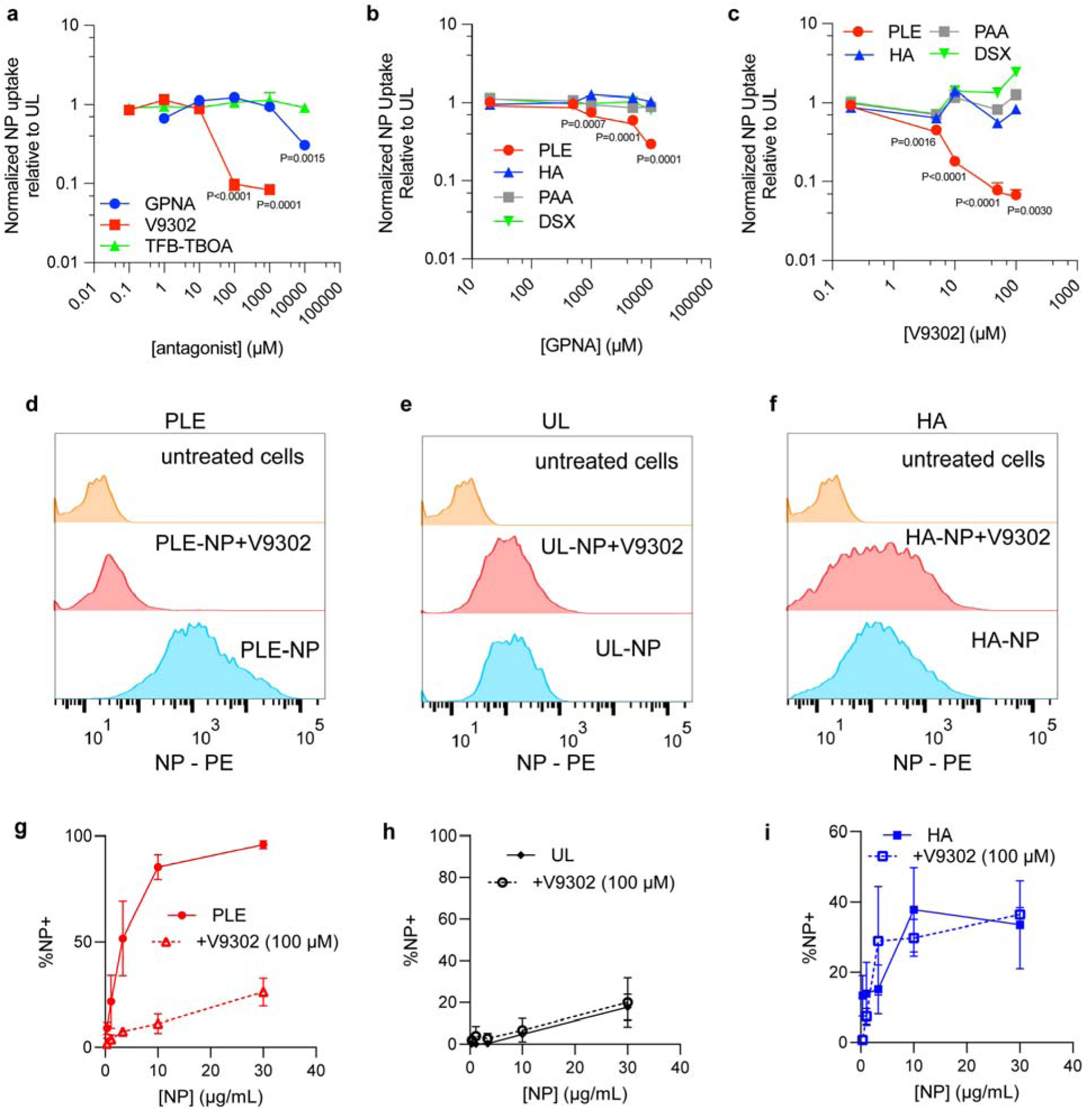
Association of PLE-coated NPs is blocked with glutamine transport inhibitors. **(a-c)** HM-1 cells were treated with varying concentrations of amino acid transport inhibitors for 15 mins before NP dosing at 50 µg/mL (**a**) or 25 µg/mL (**b-c**) per well. Four (**a**) or two (**b-c**) hours after NP dosing, wells were washed and total NP fluorescence was measured via plate reader to determine NP association. Shown are the normalized NP association of PLE-coated NPs relative to UL at different inhibitor concentrations of GPNA, TFB-TBOA and V9302 (mean ± s.d, **a**), and the normalized NP association of various outer layer LbL-coated NPs relative to UL at different inhibitor concentrations of GPNA (mean ± s.d, **b**) or V9302 (mean ± s.d, **c**). (**d-i**) HM-1 cells were treated with 100 µM of V9302 for 15 minutes prior to NP dosing at varying concentrations. Two hours after NP treatment, cells were washed and analyzed for NP association. Shown are representative NP fluorescence histograms of HM-1 cells dosed with 30 µg/mL of PLE-NPs (**d**), UL-NPs (**e**), or HA-NPs (**f**) with or without V9302 compared to untreated cells, and the percentage of NP-positive cells in PLE-NP treated (**g**), UL-treated (**h**), or HA-NP treated (**i**) HM-1s with or without V9302 across a range of NP concentrations (mean ± s.d). Statistical comparison in **a-c** was performed via two-way analysis of variance (ANOVA) with Tukey’s multiple-comparisons test comparing groups to untreated cells. Data are representative of at least two independent experiments with *n* = 3 technical replicates per group

To assess NP association on a single cell level, we preincubated HM-1 cells with a dose of V9302 shown previously to block more than 90% of glutamine uptake (100 µM^32^) and then dosed with varying concentrations of either UL-, PLE-, or HA- NPs and quantified the percentage of NP-positive HM-1 cells. While V9302 showed no impact on association for both UL- and HA-NPs, the association of the PLE-NPs was significantly negatively impacted (**Figure 2d-i**). In total, these results support that glutamine transporters play a major role in regulating the binding of PLE-NPs to cancer cells.

### Availability of SLC1A5 glutamine transporter modulates PLE-LbL NP binding to cancer cells

Both GPNA and V9302 can block glutamine import by acting on various glutamine transporters known to be overexpressed on cancer cells including SLC38A2, SLC7A5, and SLC1A5.^32,34^ Of these, both SLC7A5 and SLC1A5 were identified as associated with PLE-NP binding in our nanoPRISM analysis. SLC38A2 has limited activity in most cancer cells unless deprived of aminoacids^35^, suggesting it might be less likely to drive PLE-NP association, which we confirmed using an amino acid transporter inhibitor for SLC38A2, α-(methylamino)isobutyric acid (MeAIB), which did not affect PLE-NP association (**Extended Data Fig. 2**). While both SLC1A5 and SLC7A5 were considered potential PLE-NP binders, SLC7A5 a) depends on SLC3A2 complexation for activity which likely introduces steric hindrance, b) has substantially lower glutamine affinity than SLC1A5 and c) primarily serves for glutamine efflux, limiting its relevance as a candidate for extracellular NP binding.^12,36,37^ Thus, we focused further investigation on SLC1A5—a glutamine transporter with known overexpression in most cancer types and validated capacity for glutamate binding ^38–41^.

To further probe the relationship between PLE-NP binding and SLC1A5 expression, we utilized antibody blocking and gene knockdown experiments to observe the impact on PLE-NP association. First, to selectively block SLC1A5, we dosed HM-1 cells with antibodies directed against the transporter before dosing with either PLE-NP, UL-NPs, or HA-NPs. Anti-SLC1A5 Ab dramatically reduced PLE-NP association without affecting UL or HA-NP association (**Figure 3a-d**), confirming the specificity of PLE-NP binding to the glutamine transporter. As a cell membrane receptor control, we also evaluated the effect of an anti-CD44 Ab. Only anti- SLC1A5 and not anti-CD44 Abs (**Extended Data Fig. 3**) could dramatically reduce PLE-NP association (**Figure 3a-d**). Next, we performed a transient knockdown of SLC1A5 gene expression via siRNAs to partially reduce total SLC1A5 levels in HM-1 cells (**Figure 3e, f**). siRNA-treated HM-1 cells with reduced SLC1A5 protein expression showed a significant reduction in association with PLE-NPs while HA-NP binding was low and unaffected (**Figure 3g,h**). Together, these data suggest that PLE-NPs are capable of directly binding to SLC1A5.

**Figure 3.**
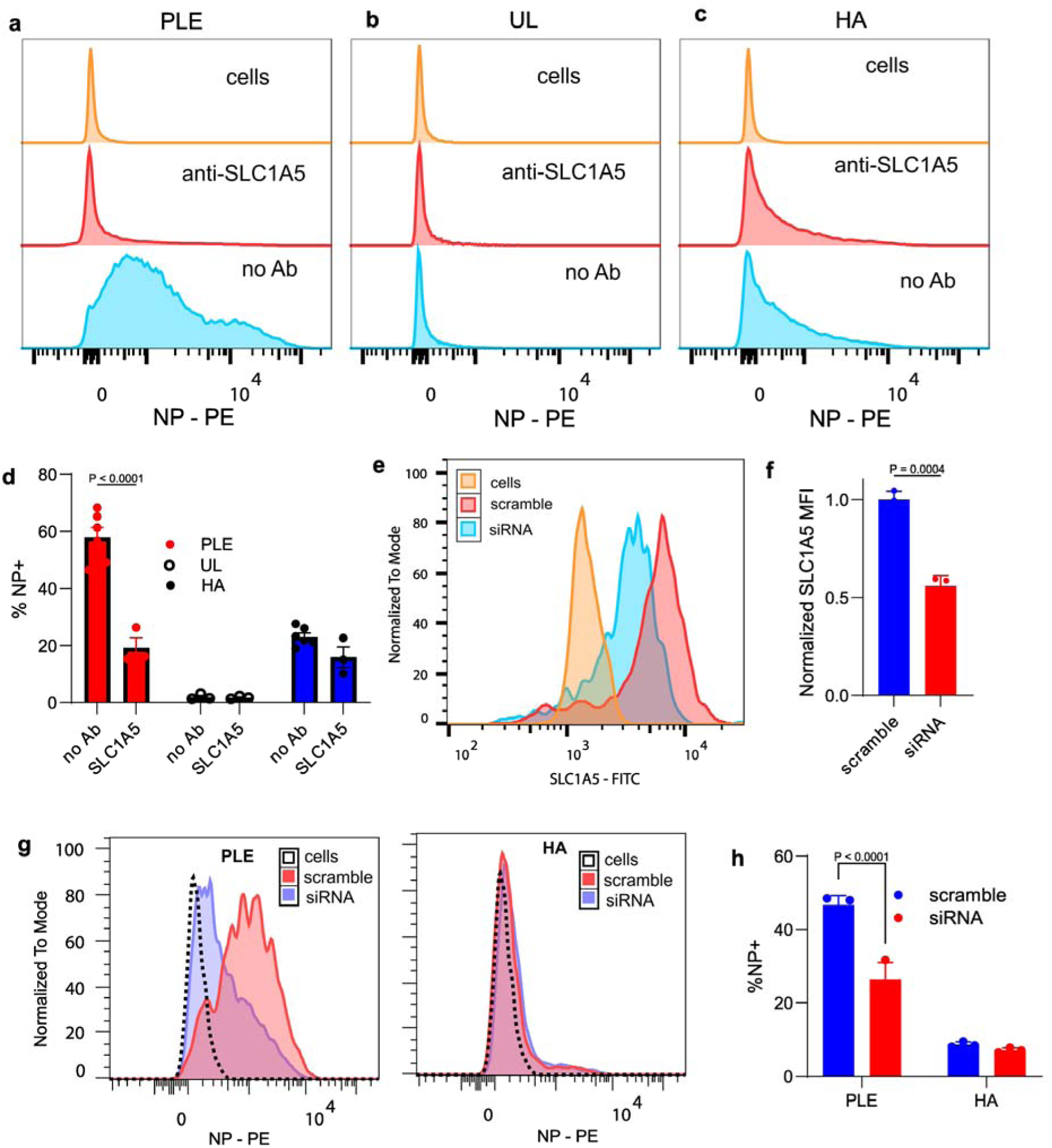
Modulation of SLC1A5 availability regulates PLE-NP binding. **(a-d)** HM-1 cells were treated with antibodies (Abs) against SLC1A5 for 1 hr. Fluorescent UL, PLE, or HA NPs (10 µg/mL) were added for 15 minutes, then cells were washed and analyzed by flow cytometry. Shown are representative histogram plots of NP fluorescence for cells incubated with PLE-NPs (**a**), UL-NPs (**b**), or HA-NPs (**c**) in the presence of anti-SLC1A5 antibody or control treatments and the percentage of NP+ cells for each treatment (mean ± s.e.m., **d**). **(e-h)** HM-1 cells were pre-treated with either anti-SLC1A5 siRNA or scramble siRNA at 200 nM for 96 hrs before dosing with 10 µg/mL of NPs for 30 min. After NP incubation, cells were washed with PBS and analyzed with flow cytometry to determine NP association. Shown are representative flow cytometry of anti-SLC1A5 Ab staining in cells treated with either scramble of anti-SLC1A5 siRNA (**e**), quantitation of median fluorescence intensity (MFI) of total anti-SLC1A5 staining (mean ± s.d, **f**), representative flow cytometry histograms of NP fluorescence of HM-1 cells with partial SLC1A5 knockdown for PLE-NP and HA-NP treatments (**g**), and the percentage of NP positive cells with partial SLC1A5 knockdown treated with PLE-NP or HA-NP (mean ± s.d., **h**). Statistical comparison in **d, h** was performed via two-way analysis of variance (ANOVA) with Tukey’s multiple-comparisons test and an unpaired t-test for **f**. Data are representative of at least two independent experiments with *n* = 3 technical replicates per group

### SLC1A5 clustering prolongs surface retention of PLE-NPs

Having discovered a binding target of PLE-NPs, we theorized that the SLC1A5 interaction might also contribute to the high cell surface retention of PLE-NPs observed in previous work where we have successfully leveraged this property to deliver interleukin-12 (IL-12) to the tumor extracellular microenvironment.^6,13,16,42^ To determine if SLC1A5 is associated with this surface retention, we utilized confocal microscopy to image HM-1 cells incubated with fluorescently-labeled IL-12-NPs with or without a PLE coating. We then stained to visualize the cellular localization of SLC1A5, revealing that UL-NPs were rapidly endocytosed with no signs of interaction with SLC1A5 (**Figure 4a, Extended Data Fig. 4a**). By contrast, PLE-NPs formed clusters on the cell surface of HM-1 cells that colocalized with accumulated SLC1A5 (**Figure 4a, Extended Data Fig. 4a** white arrows). The correlation of NPs with cell surface receptors was specific to SLC1A5 as neither CD44 nor GLUT-1 colocalized with PLE-NPs (**Figure 4a-b, Extended Data Fig. 4b-c**). We confirmed this correlation of NPs with SLC1A5 was specific to PLE NPs, as other LbL-NP coatings—including a PLR monolayer or bilayers with PLR followed by HA or polyacrylic acid (PAA)—did not show a significant correlation even when the NPs were found on the surface of cancer cells (**Figure 4c, Extended Data Fig. 4d**). PLD NPs had a low but statistically significant level of correlation with SLC1A5 when on the cell surface (**Figure 4c, Extended Data Fig. 4d**), suggesting that a polyaspartate coating may also be engaging this amino acid transporter.

**Figure 4.**
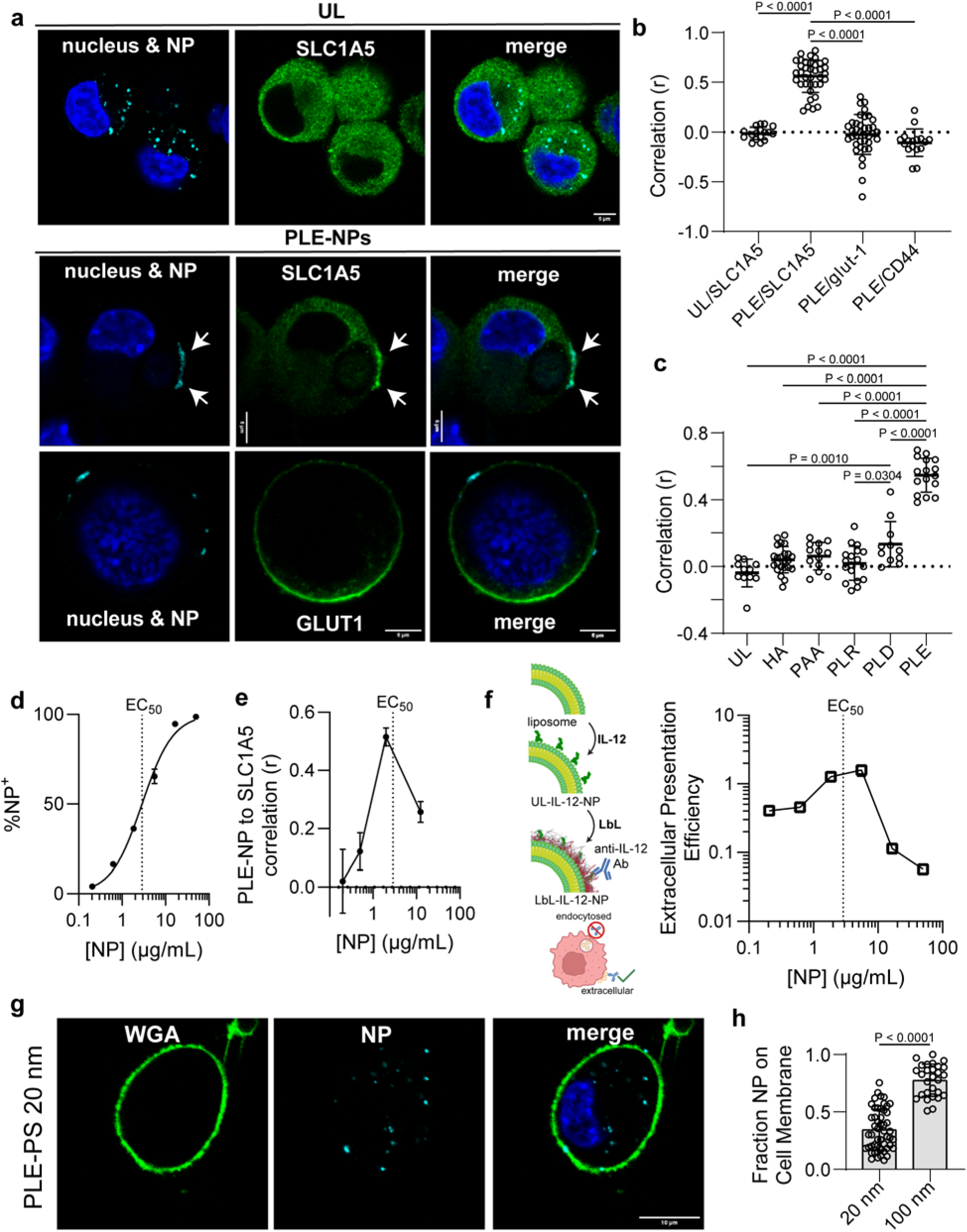
PLE-NPs colocalize with SLC1A5 transporters on the cell surface. **(a-c)** HM-1 cells were dosed with 1.5 µg/mL of NPs for 2 hrs, washed with PBS, fixed, permeabilized with saponin, and stained. Shown are representative images of HM-1 cells treated with UL-NPs or PLE-NPs and stained for SLC1A5 or GLUT-1 transporters **(a)**. A correlation analysis between NP signal and cell membrane transporter stain (mean ± s.d, **b**), and a correlation analysis between varied NP formulations and SLC1A5 staining (mean ± s.d., **c**) were conducted. Each data point represent correlation across a single cancer cell. **(d)** HM-1 cells were dosed with varying concentration of PLE-IL12 NPs for 4 hrs. Shown are the percentage of NP-positive cells for each concentration of NP dosed as determined with flow cytometry. **(e)** The same process as (**a-c**) was followed with various concentrations of LbL-NPs tested. Shown are the correlation between NP signal and SLC1A5 stain (mean ± s.d). **(f)** HM-1 cells were dosed with varying concentrations of PLE-IL-12 NPs for 4 hrs and then stained with anti-IL-12 antibody. Shown is the extracellular IL-12 presentation efficiency (the ratio between extracellular IL-12 stain to total NP uptake at 24 hrs compared to the same ratio at 4 hrs after dosing, mean ± s.d). **(g-h)** HM-1 cells were dosed with 1 µg/mL of NPs for 4 hrs, washed with PBS, fixed, and stained with Hoechst 33342 and wheat germ agglutinin (WGA) for visualization on a confocal microscope. Shown are representative HM-1 cells dosed with 20 nm PLE-PS particles (**g**) and the quantification of the fraction of NP pixels colocalized with cell membrane pixels (mean ± s.d, **h**). Statistical comparison in **b** and **c** was performed via two-way analysis of variance (ANOVA) with Tukey’s multiple-comparisons test and an unpaired t-test for **h**. Data are representative of at least two independent experiments.

While SLC1A5 itself has a low internalization rate (half-life 20-60 hrs^43,44^) that could aid in the surface retention of PLE-NPs, we hypothesized that the cell surface clustering of NPs bound to SLC1A5 could contribute to their high cell surface retention property, similar to certain galectin lattice structures.^45^ To evaluate the effect of clustering, we dosed increasing concentrations of NPs to HM-1 cells and either imaged cells via confocal microscopy or quantified NP association via flow cytometry. Near the NP EC_50_ (the NP concentration leading to half-maximal binding of particles), we could observe a clear increase in the correlation between PLE-NPs and SLC1A5 signal (**Figure 4d-e**), suggesting an increase in cluster size or clustering efficiency. However, NP doses above the EC_50_ reduced SLC1A5 clustering without affecting the total fraction of initial NP binding to the cell surface (**Figure 4d-e, Extended Data Fig. 5a-b**), likely due to the saturation of SLC1A5 receptors on the cell surface (i.e., preventing one NP from binding multiple receptors). Indeed, staining for SLC1A5 showed a lack of PLE- SLC1A5 foci formation at high doses of PLE-NPs (**Extended Data Fig. 5c**).

To determine the relationship between transporter clustering and NP internalization, we leveraged the presence of a therapeutic cargo on the NPs to probe its delivery to the cell surface or intracellularly. For this experiment, IL-12 was conjugated to fluorescent liposome surfaces, followed by LbL layering of PLR and PLE (**Figure 4f**). We have previously shown that IL-12 on LbL-coated particle surfaces is accessible to staining with an anti-IL-12 monoclonal antibody.^15^ IL-12 PLE-NPs were incubated with HM-1 cells followed by staining with anti-IL-12, and the extracellular IL-12 staining signal was compared to the total NP signal from flow cytometry at 4 and 24 hrs after dosing with various concentrations of NPs. Below the EC_50_, NPs were effectively retained on the cell surfaces with ∼50% of the IL-12 signal remaining extracellular at 24 hrs, consistent with a ∼20 hr half-life^44^ of SLC1A5 (**Figure 4f**). Strikingly, there was little to no NP uptake near the EC_50_ as the ratio of external IL-12 signal to total NP signal remained constant (value ∼1). On the other hand, NPs incubated with HM-1 cells at concentrations well above the EC_50_ showed a strong decline in the IL-12:NP signal ratio, indicating internalization and suggesting that a lack of receptor clustering at high NP doses may facilitate particle internalization.

While surface retention of PLE-NPs is desirable for the delivery of drugs targeting the extracellular space, certain drug delivery applications may benefit from rapid NP internalization. Consistent with clustering-induced surface retention of PLE-NPs mechanism presented here, we have previously shown that the addition of a co-polymer of PLE with polyethylene glycol (PEG) to the LbL film prevents surface retention of PLE-NPs on cancer cells.^7^ PEG may act to sterically inhibit cluster formation at the cell surface. We theorized that the faster internalization kinetics of small NPs may also avoid cluster assembly.^46^ Moreover, as SLC1A5 has been previously shown to colocalize with caveolin-1, small PLE-NPs may readily fit into caveolar pits which could avoid cluster formation and allow for rapid internalization.^47,48^ We thus coated carboxylated polystyrene NPs of either 20 nm or 100 nm in diameter with a PLE LbL film. As reported previously^6^, 100 nm PLE-NPs accumulated on the cell surface (**Extended Data Fig. 6**). However, when we dosed HM-1 cells with 20 nm PLE-NPs, we could readily observe internalized PLE-NPs within 4 hours after dosing (**Figure 4g-h).** In all, these results suggest that inducing the clustering of SLC1A5 transporters confers surface retention for PLE-NPs.

### Molecular modeling predicts polypeptide interactions with SLC1A5 and unique PLD- SLC1A3 interactions

SLC1A5 and other amino acid transporters evolved to transport individual amino acid monomers. To gain insight into how glutamic acid polymers such as the one in the LbL coating might interact with SLC1A5, we turned to computation modeling with AlphaFold 3.^49^ We probed the human sequences for SLC1A5 and SLC38A2 as positive and negative controls for PLE-NP interactions given the results from small molecule inhibition. To further evaluate the potential interaction with additional glutamine transporters, we also evaluated interaction with SLC7A5. Transporters were modeled interacting with small (*n* = 4) oligomers of PLE, poly-L-glutamine (PLQ), or a control polypeptide with low SLC1A5 interaction – poly-L-phenylalanine (PLF).^50^ We also included PLD in the structure prediction modeling to gain more insights into the potential differences between PLE and PLD. Previously, we demonstrated that while PLE-NPs remain on the cell surface for extended durations, PLD-NPs are gradually internalized via caveolin-mediated endocytosis following their binding to cancer cell surfaces which may be due to differences in their targets.^6^ Given the ability of Alpha Fold 3 to include ions and the requirement of sodium by these transporters, they were included in the model.^51,52^ Model prediction was performed at least twice, and the chain-pair interface predicted template modeling (ipTM) score was extracted from the top-ranked predictions. The ipTM score is used to rank a specific interface between two chains, with values above 0.8 representing confident high-quality predictions, values above 0.6 suggesting potential interactions, and below 0.3 as non-interacting.^49,53^

In the evaluation of SLC1A5, PLE and PLQ were found inside the binding pocket of the protein in its conformation facing the extracellular environment (i.e., outward-facing) with ipTM scores indicative of binding (∼0.6, **Figure 5a**). Consistent with the enrichment for glutamine transporters in its gene sets (**Figure 1e**) and the partial colocalization of PLD with SLC1A5 observed by confocal microscopy (**Figure 4c**), PLD also showed favorable interaction with SLC1A5, suggesting that PLD-NPs and PLE-NPs may both interact with similar amino acid transporters that are overexpressed in cancer cells. On the other hand, PLF had significantly reduced ipTM scores and was found outside of any known pocket (**Extended Data Fig. 7a**). Performing the same analysis on SLC38A2 with PLE, PLD, and PLQ showed that only PLQ interacted favorably (ipTM>0.6) with the binding pocket, whereas PLE and PLD had significantly lower ipTM scores and did not bind to a known pocket of SLC38A2 (**Figure 5b, Extended Data Fig. 7b)**. PLF, on the other hand, yielded a high ipTM score with SLC38A2, potentially due to the minor transport capacity of phenylalanine by this transporter.^35^ Moreover, only PLQ favorably interacted with SLC7A5, indicating a specificity of PLE and PLD to SLC1A5 (**Extended Data Fig. 7c**).

**Figure 5.**
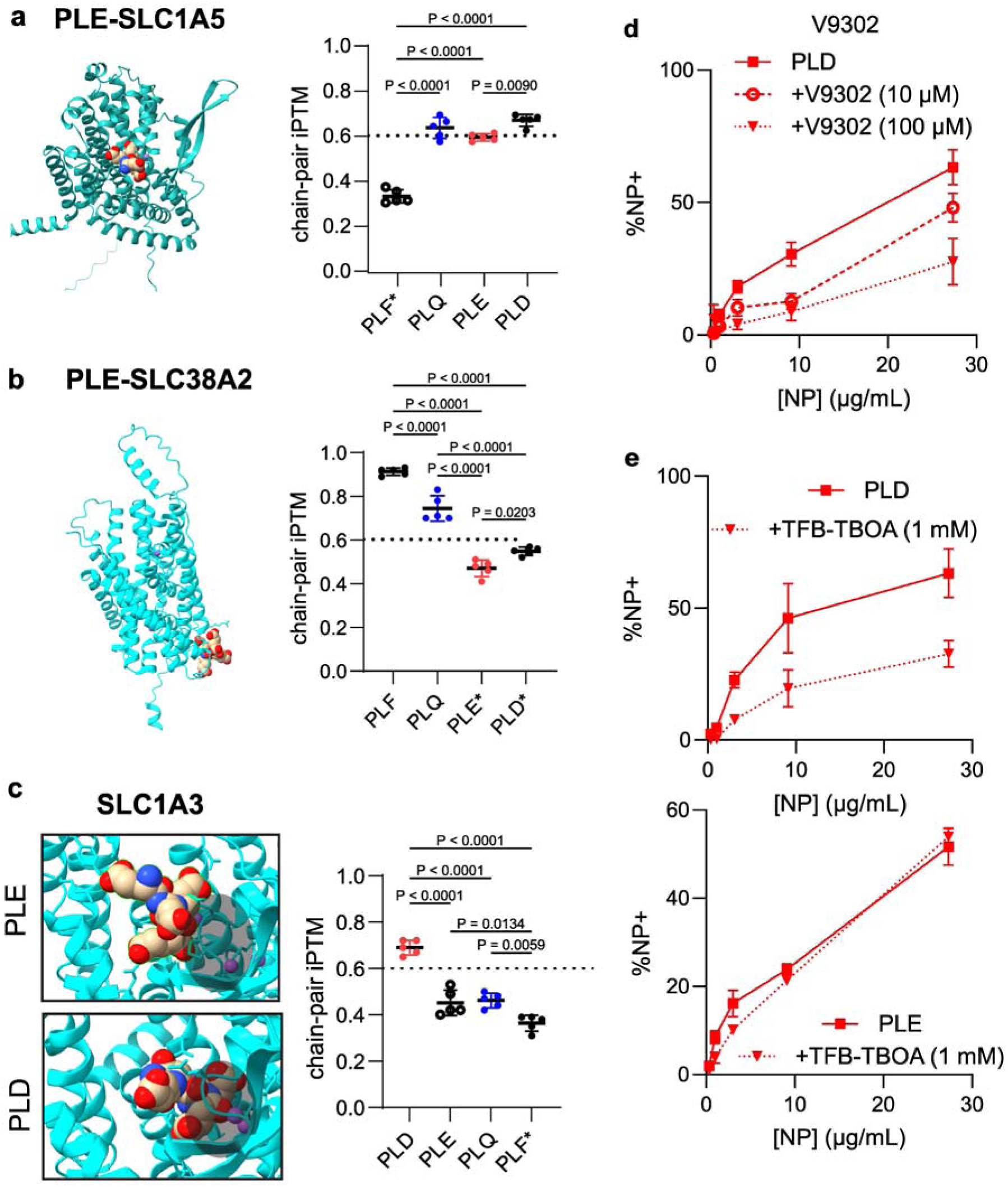
AlphaFold model predicts polymer binding to amino acid transporters. **(a)** Representative AlphaFold 3 model structure of SLC1A5 and PLE and the calculated chain-pair ipTM scores between SLC1A5 with PLF, PLQ, PLE and PLD (mean ± s.d). **(b)** Representative AlphaFold 3 model structure of SLC38A2 and PLE and the calculated chain-pair ipTM scores between SLC38A2 with PLF, PLQ, PLE and PLD (mean ± s.d). **(c)** Representative AlphaFold 3 model structure of SLC1A3 and PLE and SLC1A3 with PLD focused on the binding pocket indicated by the dark shading. AlphaFold 3 calculated chain-pair ipTM scores between SLC1A3 with PLD, PLE, PLQ, and PLF (mean ± s.d). **(d-e)** HM-1 cells were treated with either V9302 or TFB-TBOA for 15 minutes before NP dosing at varying concentrations. Two hours after NP treatment, cells were washed with PBS and analyzed with flow cytometry to measure of NP association. Shown are the percentage of NP-positive cells in PLD-treated HM-1s with V9302 (mean ± s.d, **d**), and the percentage of NP-positive cells in PLE-NP or PLD-NP treated (mean ± s.d, **e**) HM-1s with 1 mM of TFB-TBOA. (* in AlphaFold 3 iPTM scores indicates polymer not in known binding pocket of transporter). Statistical comparison in **a, b** and **c** was performed via two-way analysis of variance (ANOVA) with Tukey’s multiple-comparisons test.

Even though specialized transporters for glutamate and aspartate exist, PLE-NPs were not found to interact with these as TFB-TBOA did not inhibit PLE-NP association (**Figure 2b**). To better understand this observation, we simulated the interaction of PLE and PLD with SLC1A3, an anionic amino acid transporter that has been implicated as a major contributor to cancer progression and is overexpressed in many solid tumors.^54–57^ AlphaFold predictions showed that PLE could bind near the pocket but did not fit inside given the size of PLE’s side chain (**Figure 5c**). On the other hand, PLD was predicted to bind inside the pocket. Indeed, quantification of chain-pair ipTM scores showed that PLD had favorable binding to SLC1A3, compared to PLE’s significantly lower score which was similar to PLQ (**Figure 5c**). PLF had the lowest ipTM score and was found outside of any known binding pocket.

Based on the predicted interactions of PLD with both SLC1A5 and SLC1A3, we next sought to validate these observations experimentally. We first evaluated if the glutamine transport inhibitor V9302 found to block PLE-NP binding could also block PLD-NP association.

V9302 showed a dose-dependent inhibition of PLD-NP binding to HM-1 cells (**Figure 5d**), indicating PLD-NPs also bound to SLC1A5. We next dosed HM-1 cell with the anionic amino acid transport inhibitor – TFB-TBOA– to evaluate its effects on PLD-NPs and PLE-NP association at the single cell level via flow cytometry. Consistent with the structure modeling predictions, TFB-TBOA partially blocked PLD-NP but not PLE-NP association across NP doses (**Figure 5e**). Importantly, this ability of PLD NPs to bind to anionic amino acid transporters such as SLC1A3 explains the increased rate of PLD over PLE endocytosis since transporters of anionic amino acids have a much shorter surface half-life (<1 hr^58,59^) compared to SLC1A5 (20- 60 hrs^43,44^). Further, PLD-NP binding to SLC1A3 is consistent with the previously shown caveolin-mediated NP trafficking as SLC1A3 uptake is mediated through caveolin.^58,60^ Thus, unlike PLE, PLD coating maintains binding to anionic amino acid transporters, allowing for accelerated internalization rates.

### Expression of amino acid transporters correlates with PLE and PLD NP association across various human cancer cell lines

SLC1A5 and SLC1A3 are both overexpressed in many human cancers with expression correlating with poor prognosis.^26,41,61–64^ We theorized that the expression of these receptors may be predictive of NP binding and could be leveraged to find optimal cancers to target with these platforms. We previously screened a library of LbL-NPs including PLE- and PLD- NPs on various human ovarian cancer cell lines and primary murine non-cancerous tissues.^6^ In this study, we screened fluorescently labeled carboxylated polystyrene nanoparticles coated with either no outer layer or one of nine distinct outer layer chemistries. We then quantified cell-type-specific preferences for each outer layer using Z-score analysis.^6^

We first evaluated the correlation between PLE and PLD NPs across the cell lines as the shared binding towards SLC1A5 should yield similar association across cells. Indeed, there was a strong correlation between the increase in NP association of PLE and PLD over UL NPs in 14 human ovarian cancer cell lines and primary murine cells (**Figure 6a**). To determine if genetic markers could predict the specificity of interaction with cells, we correlated the expression levels of SLC1A5^65^ to their preference towards PLE-NPs over other NPs (PLE-NP Z-score). Except for one cell line (OVCAR4), we observed a strong relationship between PLE-NP preference and its SLC1A5 expression level (**Figure 6b**). OVCAR4 may be an outlier due to its high expression of hypoxia-related genes (**Extended Data Fig. 8**) which induce an intracellularly restricted isoform of SLC1A5.^66^

**Figure 6.**
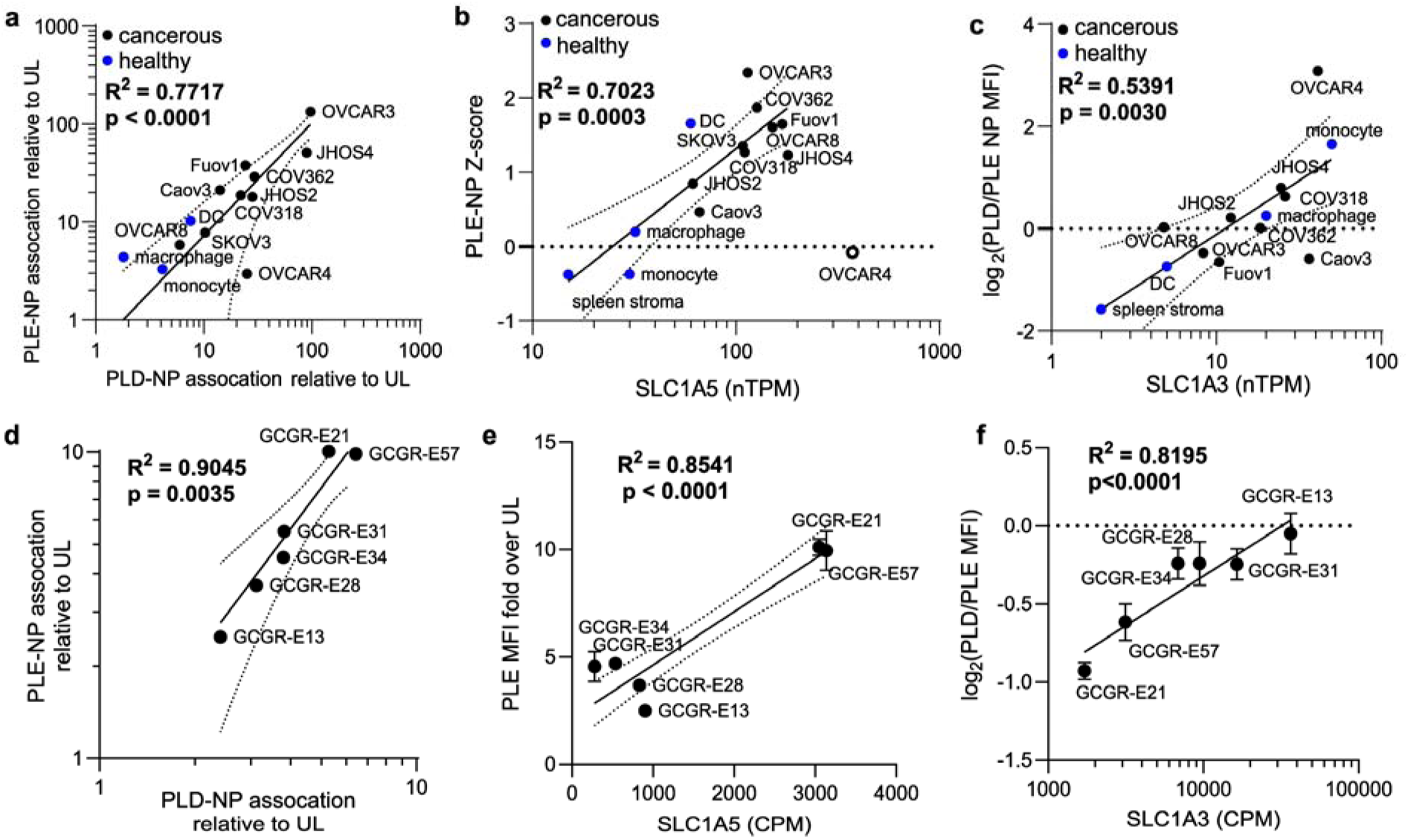
PLE-NP and PLD-NP association correlates with SLC1A5 and SLC1A3 expression. **(a-c)** Analysis of NP association with ovarian cancer cell lines and primary healthy cells with RNA expression of human cell lines derived from Protein Atlas. Shown are log-log plots of fold increase in PLE and PLD coated LbL-NPs relative to UL NPs in a library of ovarian cancer cells lines and primary healthy cells (mean, **a**), PLE-NP Z-scores from NP screen against the same cells as a function of SLC1A5 RNA expression (mean, **b**), and the ratio of NP median fluorescence intensity (MFI) of PLD-NP treated cells the MFI of PLE-treated cells as function of SLC1A3 expression (mean, **c**). **(d-f)** Same analysis as (**a-c**) but with glioblastoma cell lines (mean ± s.d.). Dashed lines represent 95% confidence interval of the curve fit. R^2^ values derived from linear fits on plots with axis as shown and P-values derived from non-zero slope test.

As expected, a similar correlation was observed between SLC1A5 and PLD-NP association (**Extended Data Fig. 9a**). However, no correlation was observed for SLC7A5 or SLC38A2, confirming the specificity of these NPs to SLC1A5 (**Extended Data Fig. 9b-e**). We next decided to explore if the expression of SLC1A3 would be predictive of the difference in binding of PLD-NPs to cells compared to PLE-NPs. We correlated the log-fold increase in PLD-NPs binding over PLE-NPs in these cells to their SLC1A3 expression and found a clear correlation (**Figure 6c**). Notably, hypoxia-related genes induce SLC1A3 expression and activity such that OVCAR4 showed the highest preference towards PLD over PLE.^67^

In addition to ovarian cancer cells, we previously showed that PLE-NPs are highly selective towards glioblastoma *in vitro* and *in vivo*.^7,29^ Consistent with these experimental observations, brain tumors are one of the cancers with the highest fold change in SLC1A5 gene expression compared to healthy tissue.^62^ Thus, to confirm the role of SLC1A5 expression levels with PLE- and PLD-NP binding, we screened their association to glioblastoma cell lines. We incubated six glioblastoma cell lines with UL-, PLE-, or PLD-NPs and quantified total NP fluorescence via flow cytometry. Similar to the ovarian cancer cell analysis, PLE- and PLD-NP delivery were found to be highly correlated (**Figure 6d)**. Moreover, there was a clear relationship between the expression levels of SLC1A5 in the tested glioblastoma cell lines, and the enhancement in NP association due to PLE (**Figure 6e**) or PLD coating (**Extended Data Fig. 9f)**. Lastly, the ratio of association with PLD- over PLE-NPs also correlated with SLC1A3 expression in these glioblastoma lines (**Figure 6f**). In all, these analyses suggest that SLC1A5 expression correlates with both PLE-NP and PLD-NP delivery while SLC1A3 expression may drive differences in cell association for PLE- and PLD-NPs.

## Conclusion

Targeting overexpressed surface markers on cancer cells to achieve specific therapeutic delivery to tumors is a major goal of drug delivery carriers. Negatively charged polypeptide LbL coatings have empirically demonstrated *in vivo* targeting of cancerous tissues and allowed for control over NP internalization rates, but their binding targets on the cancer cell surface have remained unknown. Here, we report that the high avidity presentation of anionic polypeptide coatings on LbL NPs confers targeting to amino acid transporters overexpressed in cancerous tissues. LbL coating was optimal for the proper amino acid sidechain presentation to cancer cells; whereas simply grafting polypeptide chains to NPs inhibited cell association.

To determine the surface biomarkers that act as targets for these systems, we mined prior large NP screening experiments^21^ and identified glutamine amino acid transporters within gene sets predictive of polypeptide LbL-NP association. Through small molecule inhibition, antibody blocking, and gene knockdown we identified SLC1A5 - a glutamine transporter overexpressed in many cancer types - as the major target of PLE-NPs. Further, we observed clear colocalization of PLE-NPs with SLC1A5 on the cell surface and discovered that the induced clustering of SLC1A5 enables the surface anchoring properties associated with PLE-NPs. Notably, we could design smaller PLE-NPs which, due to size, decreased clustering and led to increased NP internalization. we were also able to hypothesize that the high internalization rates of PEG-PLE copolymer LbL-NPs observed previously were due to PEG shielding of aggregate PLE interactions that lead to clustering.^7^ Combining small molecule inhibition with artificial intelligence protein structure prediction, we also show that unlike PLE-NPs, PLD-NPs interact with both SLC1A5 and SLC1A3. This dual targeting of PLD explains the difference in internalization compared to PLE NPs as SLC1A3 has higher internalization rates that proceed through caveolin-driven mechanisms as compared to SLC1A5. Moreover, we confirm these binding interactions by analyzing LbL-NP screens with ovarian cancer and glioblastoma cell lines which demonstrate a clear correlation between SLC1A5 expression with PLE- and PLD-NP association and that the preference for PLD-NPs over PLE-NPs was related to the expression of SLC1A3.

Given the increased expression of amino acid transporters in cancer cells, they have been prior targets for drug delivery. Prior work developed glutamine-grafted polymers to target amino acid transporters such as SLC1A5.^68,69^ However, as we show here, through the LbL coating, we can achieve dramatically higher affinity given its unique ability to increase surface avidity and surface presentation while enabling the design of NPs that can be retained on the cancer cell surface for extended periods. Interestingly, glutamate has been used as a control compared to glutamine with little discussion on the potential glutamate-SLC1A5 interaction which we find drives PLE-NP binding.^68,69^ Glutamate conjugation via the side chain carboxylic acid has also been employed but such an approach led to no interaction with SLC1A5 and unclear binding partners, likely due to the nature of presentation of the amino and carboxyl groups present on all amino acids.^70,71^ Grafting of glutamine residues onto branched polyethylenimine (PEI) for polyplexes has also been used^72^; however, glutamine residues are more likely to interact with other glutamine transporters such as SLC38A2 and SLC7A5. Moreover, unlike LbL-NPs, these particles lack control over internalization rates.

Beyond avidity, the high affinity of PLE and PLD towards SLC1A5 may originate from the induced protonation of their anionic residues. Polyelectrolytes tend to shift the monomer pKa due to unfavorable charge repulsion of residues in proximity. Indeed, PLE and PLD have an apparent pKa of ∼ 6.0^73^ even though the side chains of monomeric glutamate and aspartate residues have pKa of 4.3 and 3.9, respectively^74^. Interestingly, the transport and binding of glutamate to SLC1A5 is pH dependent, with greater transport at pH 6.0 than pH 7.0.^38–40^ In addition to potentially affecting charges in the binding pocket, direct proton transport is required in this system with protonated glutamate estimated to have dramatically higher affinity towards SLC1A5 compared to its charged counterpart.^39,40^ However, more work is required to validate whether the protonated glutamate is transported or if proton transport occurs via a separate path.^38^ Unfortunately, our protein structure modeling predictions do not account for local pKa factors which likely lead to underestimated interactions between PLE and PLD with SLC1A5.

To overcome this, physics-based protein interaction models that can better recapitulate the local pKa within the binding pockets of these amino acid transporters may clarify both the affinity and potential binding modes of polymers to these amino acid transporters.

Future work may aim to better understand the role of anionic amino acid transporters in mediating PLD-NP internalization. For clinical translation, it will be critical to evaluate how the surface presentation of these transporters and their expression levels on patient samples correlate with NP binding. Taken together, the data presented here provides the first demonstration that high avidity presentation of anionic polypeptides from LbL-NPs enables targeting to overexpressed amino acid transporters. These insights not only guide future LbL-NPs applications, but also now enable future rational design of next-generation polymeric coatings. Notably, the nature of these binding interactions provides insights for future clinical NP applications across multiple cancer types.

## Methods

### Materials

1,2-distearoyl-sn-glycero-3-phosphocholine (DSPC), 1,2-dioleoyl-sn-glycero-3-phosphoethanolamine-N-[4-(p-maleimidophenyl)butyramide] (sodium salt) (18:1 MPB-PE), 1-palmitoyl-2-oleoyl-sn-glycero-3-phospho-(1’-rac-glycerol) (sodium salt) (POPG), 1,2-dioleoyl-sn-glycero-3-phosphoethanolamine-N-dibenzocyclooctyl (DOPE-DBCO), 1,2-dioleoyl-3-trimethylammonium-propane (chloride salt) (DOTAP), and cholesterol were purchased from Avanti Polar Lipids. Poly-L-arginine (PLR) with a molecular weight (MW) of 9.6 kDa and poly-L-glutamic acid (PLE) with a MWs of 15 kDa (PLE_100_) or 120 kDa (PLE_800_) were purchased from Alamanda Polymers. BDP TMR azide (Lumiprobe) and BDP 630/650 azide (Lumiprobe) were conjugated to DOPE-DBCO in chloroform to generate fluorescently labeled lipids. Successful conjugation was validated via thin-layer chromatography which indicated <1% free dye. Fluorescently-labeled carboxylated polystyrene particles were purchased from ThermoFisher. V9302, α-(Methylamino)isobutyric acid (MeAIB) and TFB-TBOA, were purchased from MedChemExpress and L-γ-Glutamyl-p-nitroanilide (GPNA) was purchased from Sigma. Anti-CD44 antibody (IM7, functional grade), and anti-GLUT-1 antibody (SA0377) were purchased from Invitrogen. For anti-SLC1A5, two clones directed at extracellular epitopes were purchased - AB_2806719 from Invitrogen and AB_2878679 from Proteintech. An intracellular epitope directed anti-SLC1A5 (AB_2756720) was purchased from Alomone Labs. Anti-mouse IL-12 (clone C15.6) was purchased from Biolegend.Deionized water of the ultrapure grade was obtained through a Milli-Q water system (EMD Millipore).

### Recombinant single-chain IL-12 production

Single-chain IL-12 sequence^75^ was synthesized as a genomic block (Integrated DNA Technologies) and cloned into gWIZ expression vector (Genlantis). Plasmids were transiently transfected into Expi293 cells (ThermoFisher Scientific). After 5 days, cell culture supernatants were collected and protein was purified in an ÄKTA pure chromatography system using HiTrap HP Niquel sepharose affinity column, followed by size exclusion using Superdex 200 Increase 10/300 GL column (GE Healthcare Life Sciences). Endotoxin levels in purified protein was measured using Endosafe Nexgen-PTS system (Charles River) and assured to be <5 EU/mg protein.

### Liposome synthesis

Lipid stocks were stored at -20 °C in amber vials in chloroform. A lipid solution was prepared by mixing DSPC (25 mg/mL), cholesterol (25 mg/mL), and POPG (25 mg/mL) at a 70:24:6 mole % ratio and then forming a thin film using a rotary evaporator (Buchi). Lipid films were allowed to further dry overnight in a desiccator, then were hydrated at 0.5-1 mg/mL using deionized water and sonicated for 3-5 minutes at 65 °C then extruded (Avestin Liposofast LF-50) at 65 °C once through a 100 nm membrane (Cytiva Nuclepore) then 3 times through 50 nm membranes (Cytiva Nuclepore). Extruded liposomes were cooled in an ice bath. For fluorescence labeling of liposomes, 0.2 mol% of DSPC content was replaced by either DOPE-TMR or DOPE-630/650. After extrusion, lipid concentration was determined based on the fluorescent signal from pre-extruded lipid sample of known concentration. Cationic liposomes were made via same method but replacing POPG with DOTAP.

### IL-12 Liposome synthesis

For covalent linkage of scIL-12 to liposomes, 5 mol% of DSPC was replaced with 5% of 18:1 MPB-PE (5 mg/L) in the lipid solution prior to lipid film formation and dried in the same method as standard liposomes. For lipid film hydration, the solution pH of MPB-PE liposomes was adjusted to pH 5 with hydrochloric acid to prevent maleimide hydrolysis. Following hydration, liposomes at 0.33 mg/mL were adjusted to pH 7.0 with 10 mM HEPES followed by the addition of scIL-12 containing a terminal cysteine residue at a molar ratio of 25:1 of MPB-PE lipid to protein for at least 12 hours at 4 °C in a rotating mixer. Any remaining maleimides were quenched with a 100-fold molar excess of L-cysteine (Sigma) for 1.5 hrs on ice. Unlayered IL-12 liposomes were then purified via tangential flow filtration using a 100 kDa (mPES, Repligen) hollow fiber membrane for 7 diafiltration volumes.

For fluorescence labeling of liposomes, 0.2 mol% of DSPC content was replaced by either DOPE-TMR or DOPE-630/650. IL-12 concentrations were measured via enzyme-linked immunoassay (ELISA) (Peprotech) and lipid content was determined based on the fluorescence of the pre-extruded lipid sample of known concentration.

### Layer-by-layer (LbL) film deposition onto nanoparticles

Assembly of polyelectrolyte layers was performed by adding unlayered particles to a diH_2_O solution with 0.3-0.4 weight equivalents (wt.eq.) of PLR relative to lipid in a glass vial under sonication and incubating on ice for at least 30 min. Excess PLR polymer was purified by TFF through a 100 kDa mPES membrane (Repligen) pre-treated with a 10 mg/mL solution of free PLR. For the terminal PLE layer, purified particles coated with PLR were added to a diH_2_O solution with PLE in a glass vial under sonication at 1 wt.eq. of polymer to lipid. LbL particles were then purified by TFF on a separate 100 kDa mPES membrane (Repligen) to remove any excess PLE.

For high throughput assembly onto carboxylated polystyrene particles or varied outer-layer chemistries of IL-12 liposomes, LbL assembly was performed using a microfluidics-based method.^17^ Briefly, polymer wt.eq. were titrated to core particles based on the amount of polymer required for the onset of the plateau point of zeta potential. Then the particle solution and titrated amount of polymer were mixed using a microfluidics cartridge at equal flow rates of 10 mL/min per channel (Precision Nanosystems).

### Characterization of particle preparations

Dynamic light scattering (DLS) and zeta potential measurements were made on a Zetasizer Nano ZSP (Malvern).

### Grafted PLE-liposome assembly

To generate liposomes with grafted PLE polymers, PLE_100_ with a C-terminal azide (PLE-N3) was purchased from Alamanda Polymers. A lipid solution in chloroform with 69.8 mol% DSPC, 30 mol% cholesterol, 0.2 mol% DOPE-630/650 was allowed to dry overnight in a desiccator then suspended at 10 mg/mL in 10% MEGA-10. A separate solution with 10 mg/mL of DOPE-DBCO in 10% MEGA-10 was made and mixed with 2 molar excess of PLE-N3 and allowed to react overnight at 25 °C to generate DOPE-PLE. The two samples were mixed to generate a lipid solution containing 0.5 mol% of the DOPE-PLE and then liposome assembly was induced by diluting the sample with phosphate-buffered saline (10 mM, pH 7.4) to 0.01% MEGA-10. Samples were concentrated and purified from detergent via TFF in a 100 kDa mPES membrane.

### Fluorescent labeling of PLE polymers

PLE_100_ or PLE_800_ at 10 mg/mL was labeled by reacting with 5 molar equivalents of sulfo-cyanine3 NHS ester (Lumibrobe) in PBS adjusted to pH ∼8.5 with 0.1 M sodium bicarbonate. Excess dye was removed via extensive 0.9 wt% NaCl dialysis followed by extensive diH2O dialysis using a 3 kDa regenerated cellulose membrane (Repligen) and the purified PLE-cy3 was lyophilized until use. For assembly of LbL films with fluorescently tagged PLE, the PLE layering solution was doped with 33% of PLE-cy3.

### Cell Culture

OV2944-HM-1 cells were acquired through Riken BRC and were cultured in α-MEM supplemented with 10% FBS and 1% penicillin/streptomycin. Glioma stem cells were provided by the Pollard and Carragher Laboratories at the University of Edinburgh.^76^ Glioma cells were cultured in DMEM/HAMS-F12 supplemented with N2, B27, 10% glucose, 1% pen/strep, MEM nonessential amino acids, EGF (10 ng/mL) and FGF (10 ng/mL). Culture vessels were coated with Cultrex Laminin (R&D Systems) for 3 h at 10 μg/mL before use. Laminin was added to cell culture media at a concentration of 2 μg/mL. Media was replaced twice weekly and passaged every 5–7 days at a ratio between 1:4 and 1:6 using Accutase (Sigma). Cells were incubated in a 5% carbon dioxide humidified atmosphere at 37 °C. All cell lines were murine pathogen tested and confirmed mycoplasma negative by Lonza MycoAlert™ Mycoplasma Detection Kit.

### In vitro cellular association

HM-1 cells were plated on a tissue-culture 96-well plate at a density of 50K cells per well. The next day, wells were dosed with NPs and left for the target incubation time. For assessment of NP-associated fluorescence in a fluorescence plate reader, the supernatant was removed from the well and diluted 10X with DMSO. Cells were then washed three times with PBS and disrupted with DMSO. The fluorescence of the NPs associated with cells was then normalized to supernatant fluorescence. The relative fluorescence of each formulation was then compared to an unlayered liposome control containing the same fluorophore.

For analysis via flow cytometry, NPs were dosed at the indicated concentrations and allowed to incubate with cells at 37°C for specific number of hours in each experiment. Cells were washed with PBS then detached from the plates using 0.25% trypsin and stained with DAPI (1 µg/mL in FACS buffer, 5 min incubation) for viability assessment and fixed with 2% paraformaldehyde (30 min incubation) until analysis by flow cytometry using an LSR Fortessa (BD Biosciences). The EC_50_ of NP binding was determined based on a dose-response curve fit (Hill equation) of the percentage of NP-positive cells for each NP concentration. To estimate the number of glutamate residues per particle, we used the weight equivalents (wt. eq.) of PLE polymers required to reach the zeta potential plateau onset point (POP)^17^, together with particle size and density data. For liposomes, an average lipid molecular weight of 800 Da was assumed and the total lipid per liposomes was estimated abased on the surface area of a unilamellar liposome with a lipid headgroup size of 0.71 nm^2^. For polystyrene particles, their mass was estimated based on a sphere volume with a polystyrene density of 1 g/cm^3^.

For experiments dosed with inhibitors, 15 min prior to NP dosing, cells were treated with DMSO or inhibitor diluted 100x into cell media. For Ab blocking experiments, cells were treated with antibodies at a concentration of 5 µg/mL for 1 hr prior to NP addition. For evaluation of extracellular IL-12, after NP dosing and detachment, cells were treated with anti-IL-12 monoclonal Ab for 1 hr at 4 °C, then washed again and fixed for flow cytometry analysis.

### siRNA depletion

Knockdown of siRNAs was performed by plating HM-1 cells at 5k cells/well in a 96-well plate. The next day, cells were dosed with anti-SLC1A5 siRNA (ON-TARGETplus siRNA mouse Slc1a5, Horizon Discovery) or scramble siRNA (ON-TARGETplus Non-targeting Control Pool, Horizon Discovery) at 200 nM using Lipofectamine RNAiMAX (ThermoFihser) in Opti-MEM (ThermoFisher) medium according to manufacturer instructions. Four days after siRNA dosing, cells were washed with fresh HM-1 media and dosed with 10 µg/mL of NPs for 30 minutes prior to cell processing for flow cytometry.

### NanoPrism dataset analysis

We rank-ordered PRISM cells based on their weighted average NP-cell association after incubation with NPs for 24 hrs as measured in the previously published NanoPrism screen.^21^ This metric is indicative of NP association with a cancer cell.^21^ The top 100 and bottom 100 cell lines were then compared for differential gene expression within each NP group (i.e., top 100 PLE-NP associated lines compared to the bottom 100 PLE-NP associated lines), and the log-fold change in gene expression and p-value for each gene were generated using the OmicsExpressionProteinCodingGenesTPMLogp1 dataset from DepMap (Public 25Q2) and DESeq2 package in R.^77^ For each NP formulation evaluated, this information was used to generate a rank ordered list (calculated as sign(log_2_(foldchange))*-log_10_(p_value_)) to then perform gene set enrichment analysis (GSEA) with the Hallmark Gene Sets.^78,79^ The normalized enrichment scores were used for visualization (FDR q value < 0.05), and only Hallmark Gene Sets with significant enrichment in at least one of the three NP groups were shown.

### Confocal Microscopy

For confocal imaging, 8-well chambered coverglasses (Nunc Lab-Tek II, Thermo Scientific) were coated with rat tail collagen type I (Sigma-Aldrich) per the manufacturer’s instructions. HM-1 cells were plated into the wells at a density of 10K/well and left to adhere overnight prior to NP treatment. After the desired incubation time with NPs, cells were washed 3x with PBS. After washing, cells were fixed in 4% paraformaldehyde for 15 minutes then washed (3x with PBS) and stained. For cell membrane staining, wheat germ agglutinin (WGA) conjugated to Alexa Fluor488 (Invitrogen) was used. For cell membrane receptor staining, cells were permeabilized for 10 min with 0.1% saponin (Sigma), washed (3x PBS), and then incubated with primary antibodies directed at intracellular epitopes of SLC1A5 or GLUT-1 in 0.5% BSA for at least 3 hrs at 25 °C. Cells were then washed (3x PBS) and stained with secondary anti-rabbit IgG conjugated to Alexa Fluor488 (Invitrogen) for 30 minutes (for anti-CD44 staining, anti-rat IgG AlexaFluor488 was used). Hoechst 33342 (Thermo Scientific) nuclear staining was included in all preparations following manufacturer’s instructions. Images were analyzed using ImageJ. Slides were imaged on an Olympus FV1200 Laser Scanning Confocal Microscope.

Confocal images of HM-1 cells at a 60x magnification were analyzed on ImageJ using the Correlation Threshold function to determine the correlation between receptor staining and NP fluorescence. Each point represents a single cell.

For evaluation of the percentage of NP signal colocalized on the surface of HM-1 cells, an RO1 on the cell was used and the same function was applied to determine the Mander’s Coefficient between the NP signal and the wheat germ agglutinin cell membrane stain.

### AlphaFold 3 Artificial Intelligence Modeling

The AlphaFold Server running AlphaFold 3 was used to predict the interaction of four residue-long polypeptides with the human protein sequences of amino acid transporters. Four sodium ions were included in the model. The highest-ranking model predicted for each job in which the polypeptide was docket to the outward-facing binding pocket was used. The highest overall ranking prediction was used if none of the five predictions yielded attachment to the binding pockets. Each amino acid transporter pair was modeled five times and the output inter-chain ipTM between the polypeptide and the transporter was extracted from the model result.

### Ovarian cancer cell line NP association screen

Previously published data^6^ on the association of fluorescently labeled unlayered and layered carboxy-modified latex nanoparticles (NPs) with 10 human ovarian cancer cell lines and 4 healthy primary murine cell types were used for subsequent analyses. To avoid confounding effects from biologically quiescent populations without appreciable endocytic or surface transporter activity lymphocytes (B- and T-cells) were not included. In the prior study, cells were exposed to NPs for 24 hours, and NP association was quantified by flow cytometry. Z-scores were calculated for each NP formulation, reflecting the relative preference of each cell line for a given NP based on outer-layer polymer identity. In the present work, we assessed the correlation between target gene expression and the fold change in cell-associated median fluorescence intensity (MFI) of PLE- and PLD-coated NPs relative to unlayered NPs. In addition, previously reported Z-scores and the difference in MFI between PLD- and PLE-coated NPs were correlated with gene expression profiles from human ovarian cancer cell lines and healthy primary cells, using single-cell RNA sequencing data from the Human Protein Atlas.^80^

### Glioblastoma cell line NP association screen

Glioblastoma cell lines (GCGR-E21, GCGR-E57, GCGR-E34, GCGR-E31, GCGR-E28, GCGR-E13), were seeded on 96-well plates at a density of 15k cells per well in 100 μL of the appropriate culture media and allowed to adhere overnight. Fluorescent LbL-NPs and UL-NPs generated as described previously were dosed at 5 µg/mL.^7^ After 24 hrs of incubation, cells were washed three times with PBS and detached with 25 μL of trypsin-EDTA or Accutase. 200 μL of FACS buffer (PBS with 1% bovine serum albumin and 1 mM EDTA) with 1 μg/mL propidium iodide (Thermo Fisher) was used to quench the dissociation, and cells were pipetted vigorously to achieve a single-cell suspension. Cells were transferred to a new 96-well plate without any laminin or PLL coating, and samples were analyzed using a BD LSR II Flow Cytometer. Gene expression data was provided by the Glioma Cellular Genetics Resource

### Statistical Analysis

GraphPad PRISM 10 was used to perform statistical analyses. Comparisons between two groups was performed via unpaired t-tests. For multiple groups or multiple variable analysis, one-way, or two-way ANOVAs were used with Tukey’s posthoc correction for time-based analysis or Sidak posthoc for other ANOVA analysis.

## Acknowledgments

We thank the Koch Institute Swanson Biotechnology Center for technical support. We also thank Gillian M. Morrison, Steven M. Pollard, and The Glioma Cellular Genetics Resource (; https://github.com/GCGR) for providing glioblastoma cell lines and associated expression data.

## Data availability

Raw data are available from the corresponding author upon request.

