## Extended Data for "Surface avidity of anionic polypeptide coatings target nanoparticles to cancer-associated amino acid transporters"

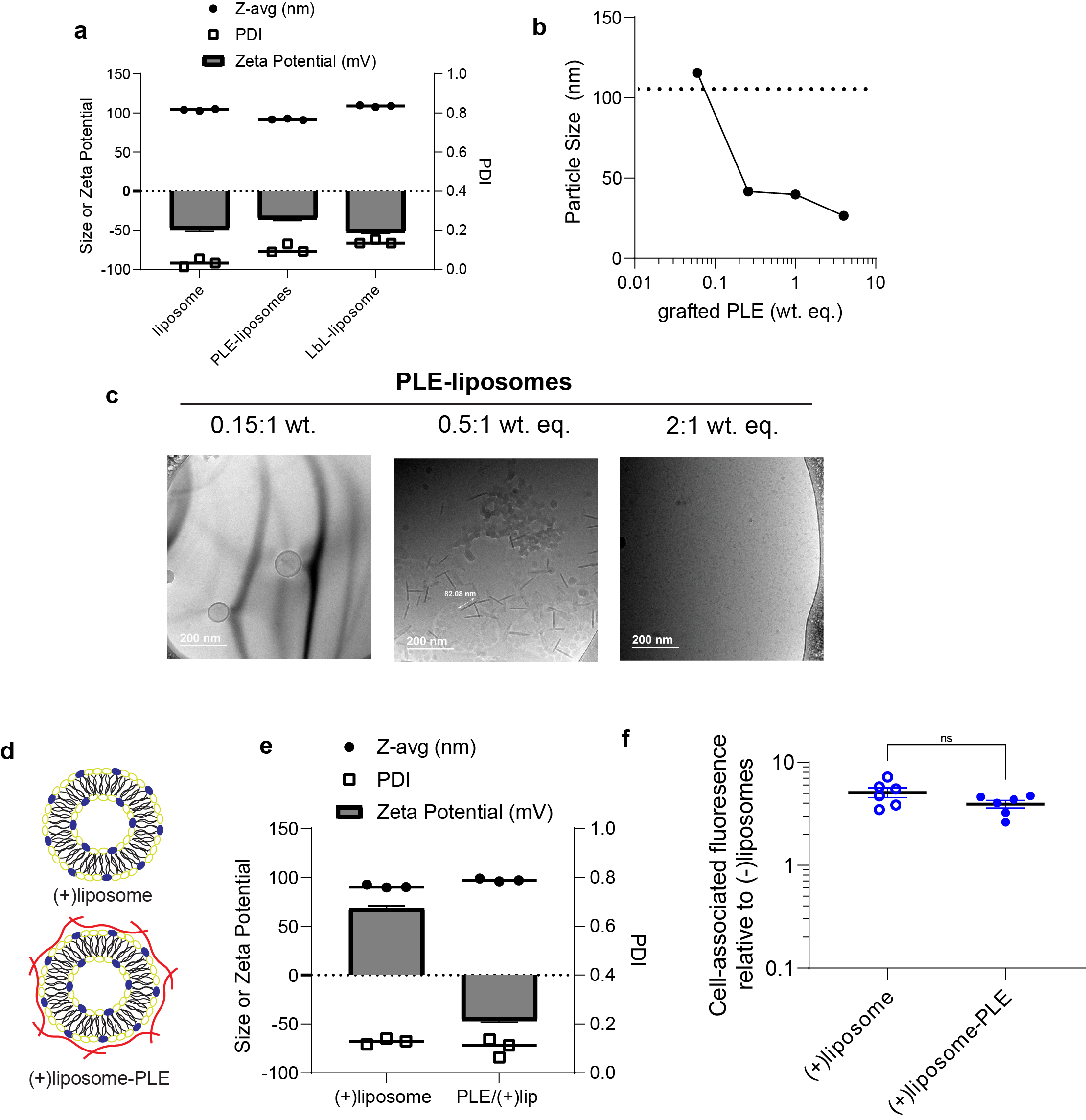

**Extended Data Fig. 1. Characterization of varied NP surface PLE presentation.** (**a**) Size (Z-avg), polydispersity index (PDI), and zeta potential (mean ± s.d.) of standard anionic unlayered liposomes, PLE-grafted liposomes (PLE-liposomes), and anionic liposomes coated with bilayer of PLE and PLR (LbL-liposomes). (**b**) Particle size upon assembly of PLE-liposomes with varied wt.eq. of PLE grafted to lipid. (**c**) Representative cryo-TEM micrograph of assembled NPs with increasing weight equivalents of grafted PLE. (**d-e**) Cationic liposomes were generated and layered with PLE and evaluated for association with HM-1 cells. Shown are the schematic (**d**) and size (Z-avg), PDI, and zeta potential (**e**). (**f**) Fluorescently labeled NPs dosed at 1 µg/mL in HM-1 cells, washed after 4 hr incubation and NP fluorescence associated with cells then measured on plate reader. Statistical comparison in **f** was performed via unpaired t-test.

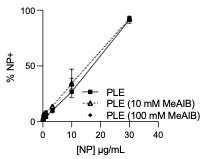

**Extended Data Fig. 2. System A amino acid transport inhibitor MeAIB does not impair PLE-NP binding.** HM-1 cells were plated in 96 well plates at 50 k cells/well and left to adhere overnight. Cells were then treated with 10 or 100 mM of MeAIB for 15 minutes prior to NP dosing at varying concentrations. Two hours after NP treatment, cells were washed with PBS, and suspended for flow cytometry analysis of NP uptake. Shown are the percentage of NP+ cells at each concentration of NP dosed.

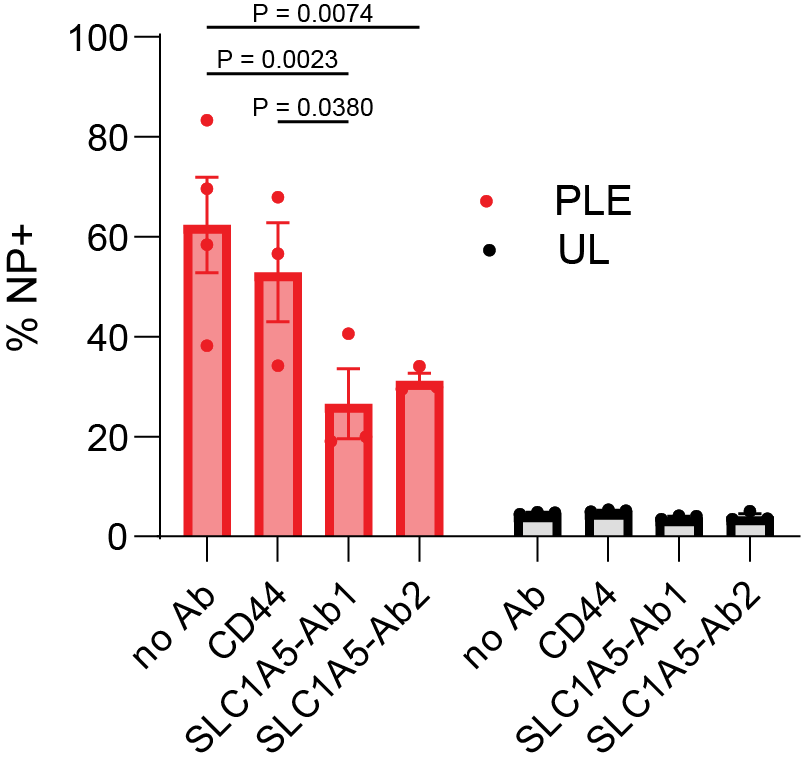

**Extended Data Fig. 3. Ab blockade of PLE-NP association is specific to anti-SLC1A5.** HM-1 cells were treated with antibodies (Abs) against CD44 or two clones against SLC1A5 for 1 hr. Fluorescent UL, PLE, or HA NPs (10 µg/mL) were added for 15 minutes, then cells were washed and analyzed by flow cytometry. Shown are the percentage of NP+ cells for each treatment

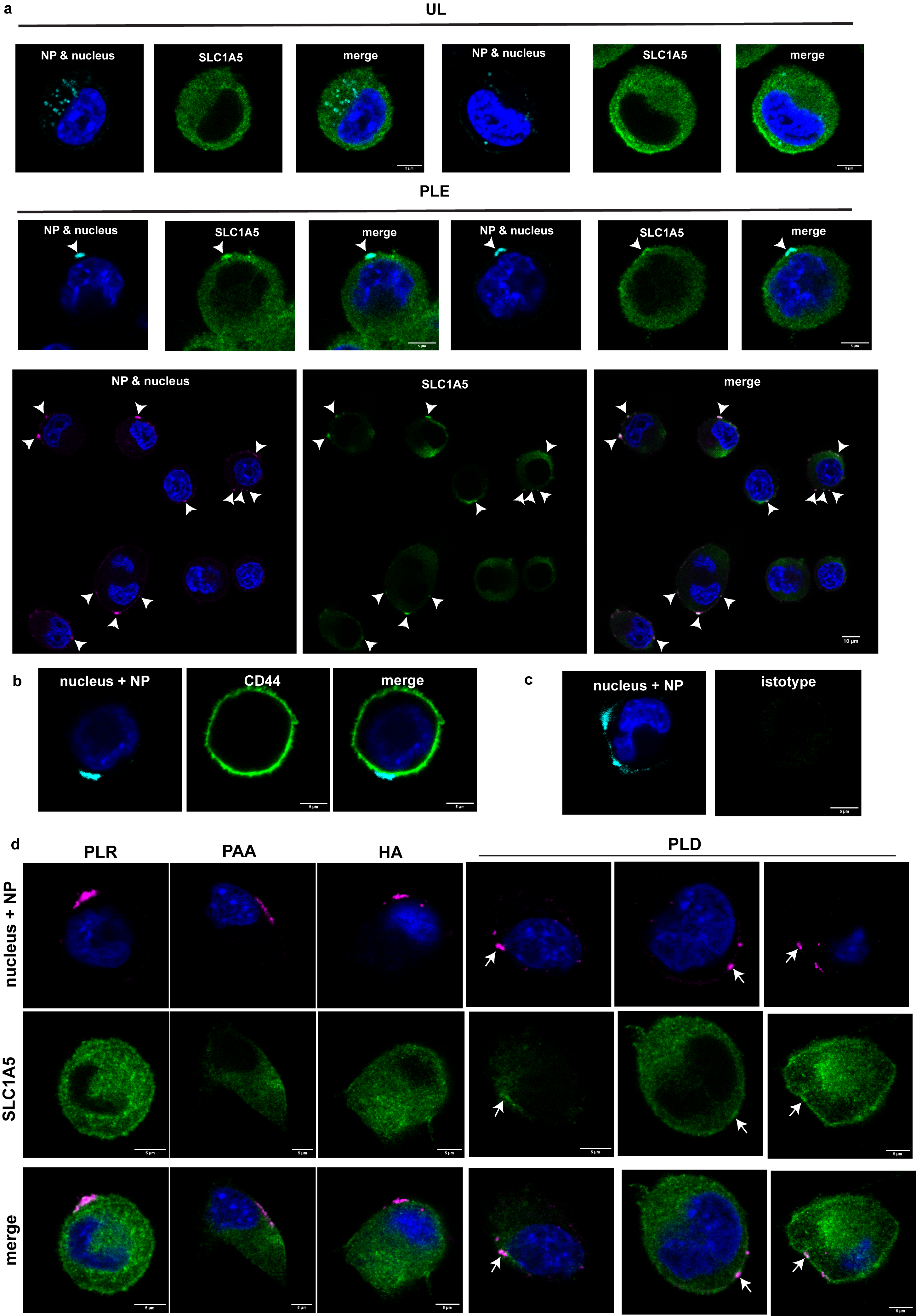

**Extended Data Fig. 4. PLE coating is required for high colocalization of SLC1A5 with LbL-NPs.** (a-d) HM-1 cells were plated in 8-well glass chamber slides at 10 k cells/well and left to adhere overnight. Cells were dosed with 1.5 µg/mL of NPs for 2 hrs. After NP treatment, cells were washed with PBS, fixed with PFA, and then rapidly permeabilized with saponin. Cells were then treated with primary antibodies for 2 hours followed by secondary antibodies for 30 minutes. Shown are HM-1 cells treated with either PLE-NP or UL-NPs and stained with anti-SLC1A5 (a), PLE-NPs and stained with an anti-CD44 Ab (b) or isotype control (c), and HM-1 cells treated with various outer layer LbL-NPs and stained with anti-SLC1A5 Abs (d).

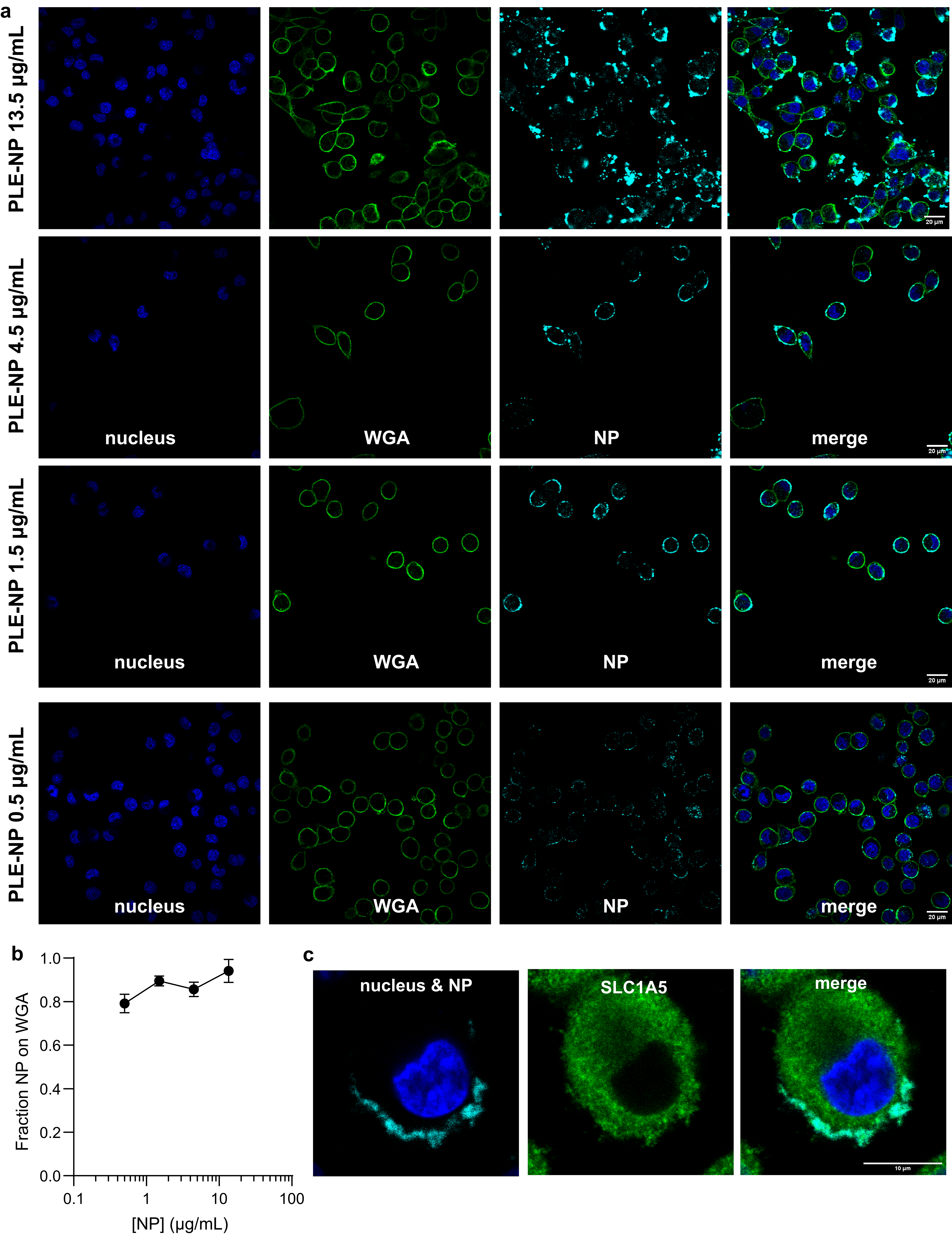

**Extended Data Fig. 5. PLE-NPs associate primarily on the cell membrane.** (a-b) HM-1 cells were plated in 8-well glass chamber slides at 10 k cells/well and left to adhere overnight. Cells were dosed with varying concentrations of NPs for 4 hrs. After NP treatment, cells were washed with PBS, fixed with PFA, and then stained with Hoechst 33342 and wheat germ agglutinin (WGA) and visualized on a confocal microscope. Shown are representative confocal images of HM-1 cells treated with various concentrations of PLE-NPs (a) and quantification of the fraction of NP pixel colocalized with cell membrane pixels (b). (c) HM-1 cells were plated in 8-well glass chamber slides at 10 k cells/well and left to adhere overnight. Cells were dosed with 13.5 µg/mL of NPs for 2 hrs. After NP treatment, cells were washed with PBS, fixed with PFA, and then rapidly permeabilized with saponin. Cells were then treated with primary antibodies for 2 hours followed by secondary antibodies for 30 minutes.

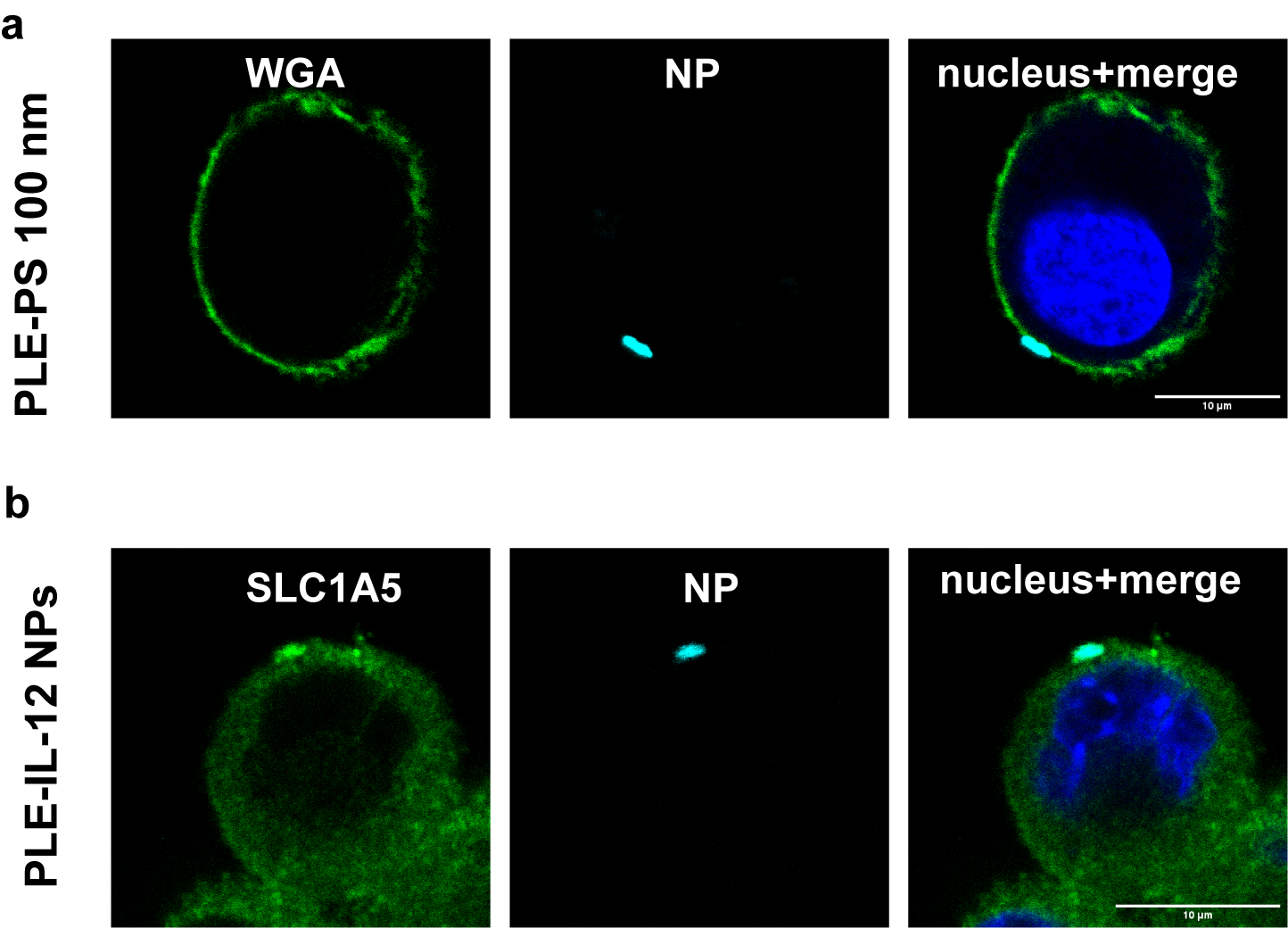

**Extended Data Fig. 6. 100 nm polystyrene PLE-NPs are retained at the cell membrane.** HM-1 cells were plated in 8-well glass chamber slides at 10 k cells/well and left to adhere overnight. Cells were dosed with 1 µg/mL of NPs for 4 hrs. After NP treatment, cells were washed with PBS, fixed with PFA, and then stained with Hoechst 33342 and wheat germ agglutinin (WGA) and visualized on a confocal microscope. Shown is a representative HM-1 cell dosed with 100 nm PLE-PS particles.

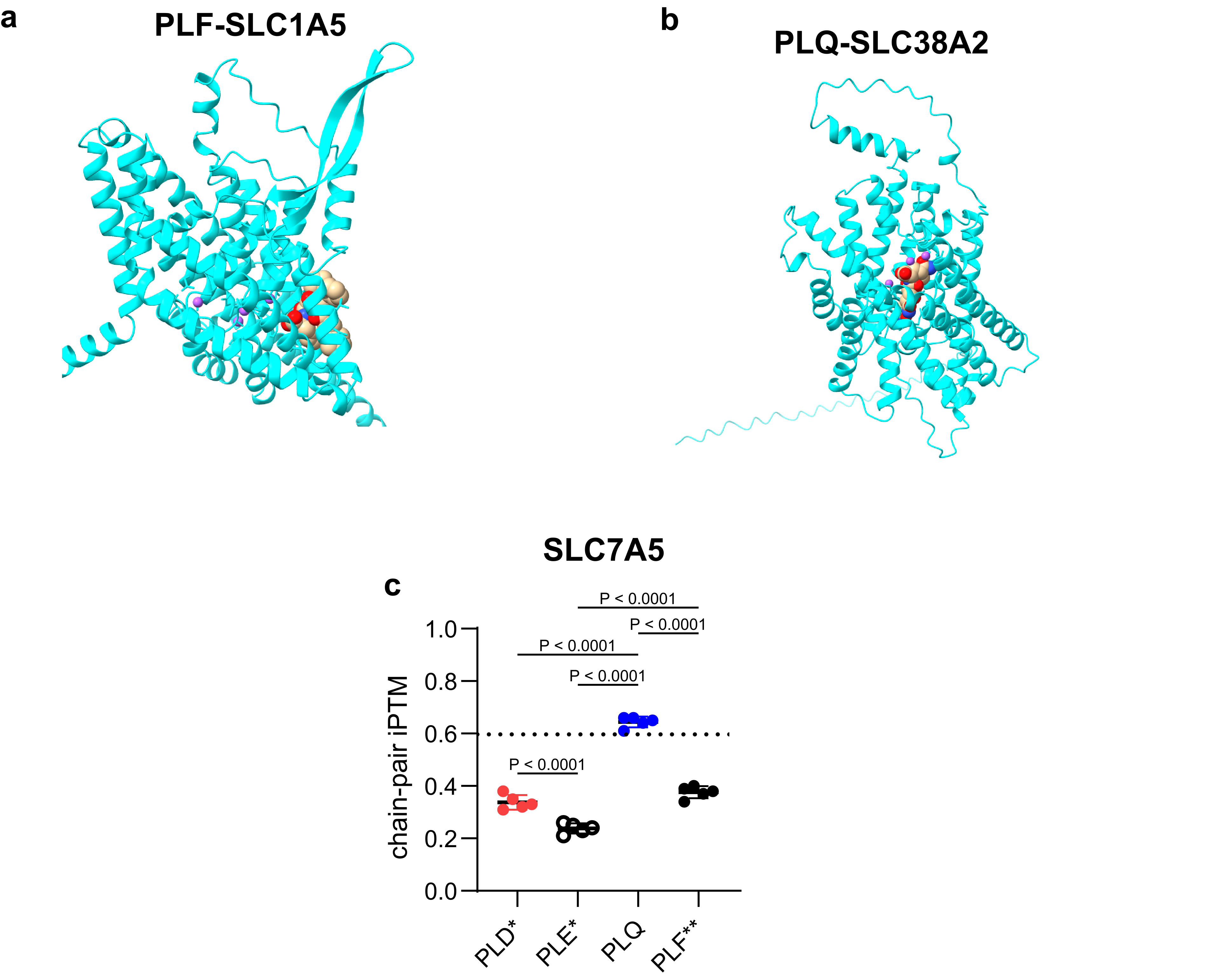

**Extended Data Fig. 7. Modeling predictions.** (a) Representative AlphaFold 3 model structure of SLC1A5 and PLQ bound to the outward or inward orientation of SLC1A5. (b) Representative AlphaFold 3 model structure of SLC1A5 and PLF binding to a transmembrane region. (c) AlphaFold3 chain pair iPTM scores of modeled PLD, PLE, PLQ, and PLF with SLC7A5 (* indicates polymer not in known binding pocket of transporter).

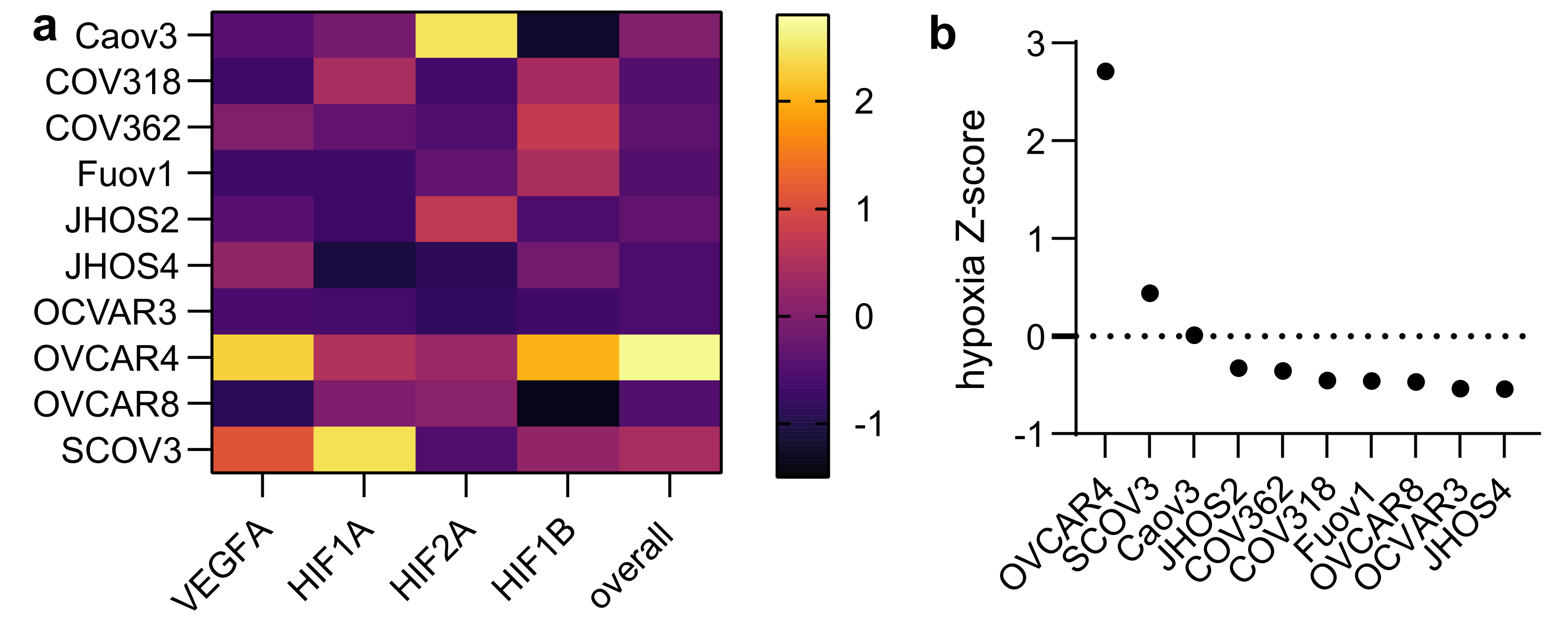

**Extended Data Fig. 8. Analysis of hypoxia-related gene expression in ovarian cancer cell lines.** (a) Heat map of Z-score for each gene across the cell lines and the “overall” hypoxia metric determined by the product of gene expression for each gene. (b) Z-score of the overall hypoxia expression levels. Expression levels extracted from the Protein Atlas.

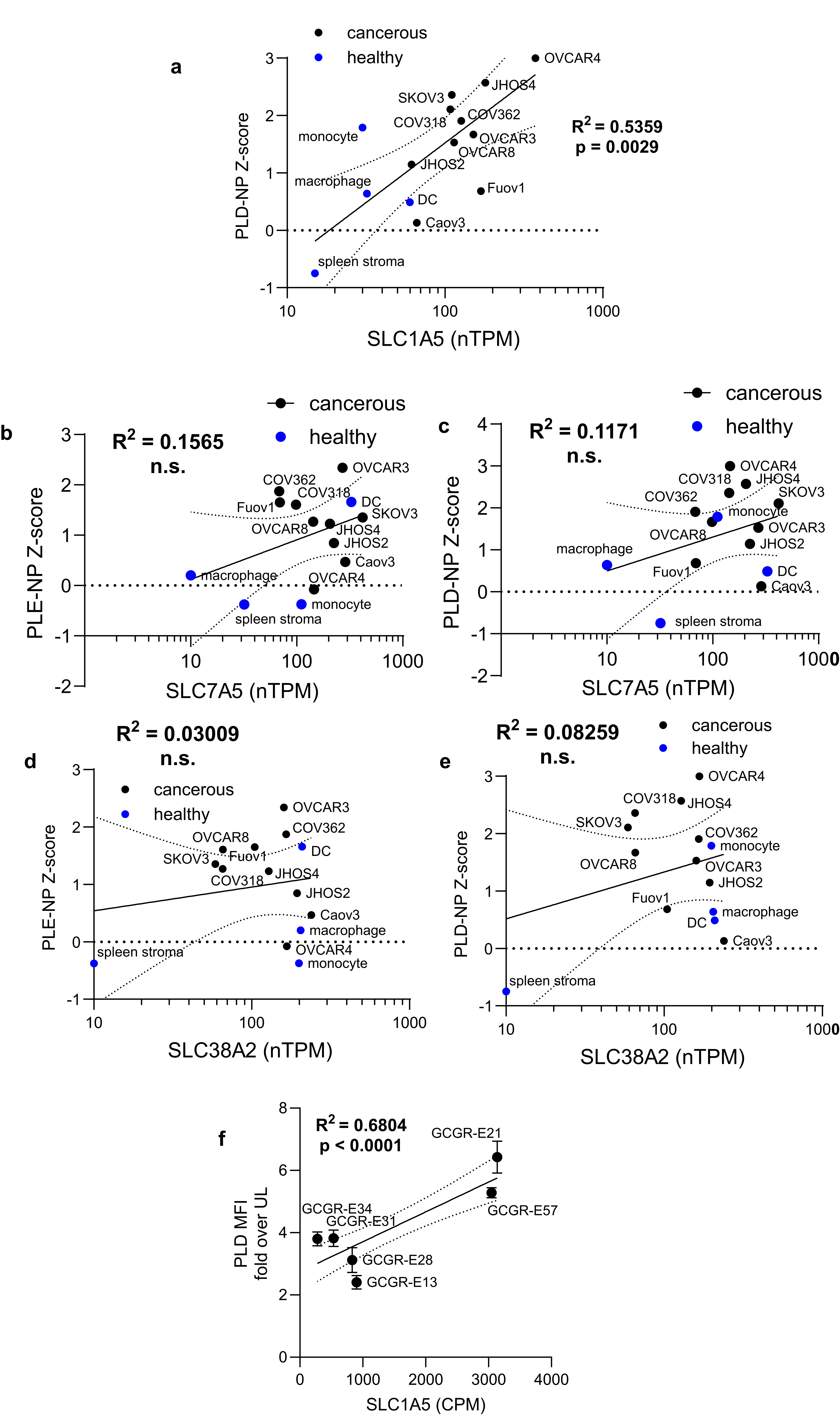

**Extended Data Fig. 9.** (a-e) Analysis of NP association with ovarian cancer cell lines and primary healthy cells with RNA expression of human cell lines derived from Protein Atlas and R^2^ found via linear fit with p-value from non-zero slope test. Dashed lines represent 95% confidence interval of the curve fit. (a) Analysis of PLD-NP Z-scores from NP screen against the same cells as a function of SLC1A5 RNA expression. Analysis of PLE (b, d) and PLD (c, e) Z-scores from NP screen against the same cells as a function of SLC7A5 and SLC38A2 RNA expression. Same analysis as (a-e) but with glioblastoma cell lines and PLD MFI over UL as a function of SLC1A5 mRNA expression.
